# Breast Cancers That Disseminate to Bone Marrow Acquire Aggressive Phenotypes through CX43-related Tumor-Stroma Tunnels

**DOI:** 10.1101/2023.03.18.533175

**Authors:** Saptarshi Sinha, Brennan W. Callow, Alex P. Farfel, Suchismita Roy, Siyi Chen, Shrila Rajendran, Johanna M. Buschhaus, Celia R. Espinoza, Kathryn E. Luker, Pradipta Ghosh, Gary D. Luker

## Abstract

Estrogen receptor-positive (ER+) breast cancer commonly disseminates to bone marrow, where interactions with mesenchymal stromal cells (MSCs) shape disease trajectory. We modeled these interactions with tumor-MSC co-cultures and used an integrated transcriptome-proteome-network-analyses workflow to identify a comprehensive catalog of contact-induced changes. Conditioned media from MSCs failed to recapitulate genes and proteins, some borrowed and others tumor-intrinsic, induced in cancer cells by direct contact. Protein-protein interaction networks revealed the rich connectome between ‘borrowed’ and ‘intrinsic’ components. Bioinformatics prioritized one of the ‘borrowed’ components, *CCDC88A*/GIV, a multi-modular metastasis-related protein that has recently been implicated in driving a hallmark of cancer, growth signaling autonomy. MSCs transferred GIV protein to ER+ breast cancer cells (that lack GIV) through tunnelling nanotubes via connexin (Cx)43-facilitated intercellular transport. Reinstating GIV alone in GIV-negative breast cancer cells reproduced ∼20% of both the ‘borrowed’ and the ‘intrinsic’ gene induction patterns from contact co-cultures; conferred resistance to anti-estrogen drugs; and enhanced tumor dissemination. Findings provide a multiomic insight into MSC→tumor cell intercellular transport and validate how transport of one such candidate, GIV, from the haves (MSCs) to have-nots (ER+ breast cancer) orchestrates aggressive disease states.

## Introduction

Estrogen receptor-positive (ER+) breast cancer, the most common subtype, preferentially disseminates to bone and bone marrow (1, 2). Bone metastases remain incurable and cause debilitating symptoms, e.g., pain, fractures, and life-threatening hypercalcemia (3). Bone marrow harbors disseminated tumor cells (DTCs) in breast cancer. DTCs may persist in a clinically occult state for decades before proliferating and potentially traveling to other organs to produce delayed recurrences in ∼40% of patients with ER+ breast cancer (4). Recurrent disease is more aggressive, relatively resistant to treatment, and currently incurable. Little progress has been made in treating breast cancer that has already disseminated to bone (5), largely because we lack a comprehensive understanding of molecular mechanisms that make ER+ breast cancer cells more aggressive in the bone marrow niche (6).

Interactions with stromal cells in the bone marrow environment are proposed to shape key functions of breast cancer cells that determine disease progression and outcomes (7). Mesenchymal stem cells (MSCs), a multipotent cell type that contributes to formation of bone, fat, and cartilage, are a major stromal cell type driving aggressive phenotypes of disseminated ER+ breast cancer cells in bone marrow. MSCs regulate ER+ breast cancer cells through mechanisms such as secreted cytokines and direct intercellular interactions (8). Work by us and others has established that direct interactions with MSCs, rather than soluble molecules, drive changes in gene expression and metabolism that promote cancer stem-like cell states, resistance to anti-estrogen drugs, and metastasis in ER+ breast cancer (9–11). Breast cancer cells and MSCs can establish channels of intercellular communication that transport molecules and organelles. Two major structures for intercellular communication include gap junctions and tunneling nanotubes (TNTs), both of which share connexin 43 (Cx43) as an essential molecular player. Gap junctions are intercellular channels comprised of various subsets of connexin proteins (12). Lymph node and systemic metastases in breast cancer and other malignancies upregulate gap junctions and Cx43, suggesting these structures contribute to essential steps in the metastatic cascade (13). TNTs are actin-rich long, thin tubes that may form between cells of the same or different types, thereby allowing long-range communication via the exchange of materials (14, 15). TNTs serve as ‘exchange freeways’ for a broad range of intracellular contents, including microRNAs, proteins, and organelles. Such exchanges, between tumor cells or between tumor and stromal or immune cells (8, 16, 17) shape drug resistance of cancer cells; regulate proliferation; and promote metastasis.

Despite these insights, only a limited number of molecules transferred from MSCs via gap junctions or TNTs have been implicated in aggressive phenotypes of breast cancer cells, whereas the identity of most remains unknown. We tackle this knowledge gap by performing a comprehensive multiomic network analysis to identify molecules and biologic processes regulated by direct contact between bone marrow MSCs and ER+ breast cancer cells. We identified genes and proteins that cancer cells acquire from MSCs through direct transfer (“borrowed” components) and additional molecules induced because of such transfer (“intrinsic components”). We used bioinformatic approaches to explore the implications of these gene expression changes on patient outcome and subsequently prioritized one ‘borrowed’ gene/protein (*CCDC88A/*GIV) to explore further and validate through a series of *in vitro* and *in vivo* studies. Our findings establish a comprehensive resource for changes induced in ER+ breast cancer cells by contact with bone marrow MSCs and pinpoint GIV as a promising target to overcome aggressive phenotypes acquired by breast cancer cells in bone marrow.

## Results

### Multiomic analysis of close contact between ER+ breast cancer cells and MSCs

DTCs in bone marrow displace hematopoietic stem cells from supportive niches (18), where they establish direct contact with various subsets of MSCs and eventually gain aggressiveness. To identify how MSCs may impact ER+ breast cancer cells through direct contact, we used a 2D co-culture model combining MCF7 or T47D human ER+ breast cancer cells with human HS5 or HS27a MSCs in a 1:9 ratio (**Fig 1A**). After 3 days of co-culture in medium with low serum and physiologic concentrations of glucose, we recovered cancer cells using EpCAM immunomagnetic beads, capitalizing on expression of EpCAM exclusively on tumor cells of epithelial origin. We previously showed that such EpCAM-based immunoisolation has negligible MSC contamination(19) and confirmed through rigorous characterization studies showing that tumor cells subjected to such contact culture gain advantageous features such as metabolic plasticity(20), resistance to estrogen-targeted therapies, and enhanced survival of disseminated tumor cells early in the metastatic process(19, 21). To control for effects of soluble molecules produced by MSCs, we cultured MCF7 or T47D cells for 3 days in conditioned medium from MSCs (**Fig 1A**). We used monocultures of MSCs and tumor cells as additional comparator groups. We analyzed all samples by bulk RNA sequencing; we also analyzed mono- and contact cultured MCF7 cells by TMT proteomics.

**Figure 1.**
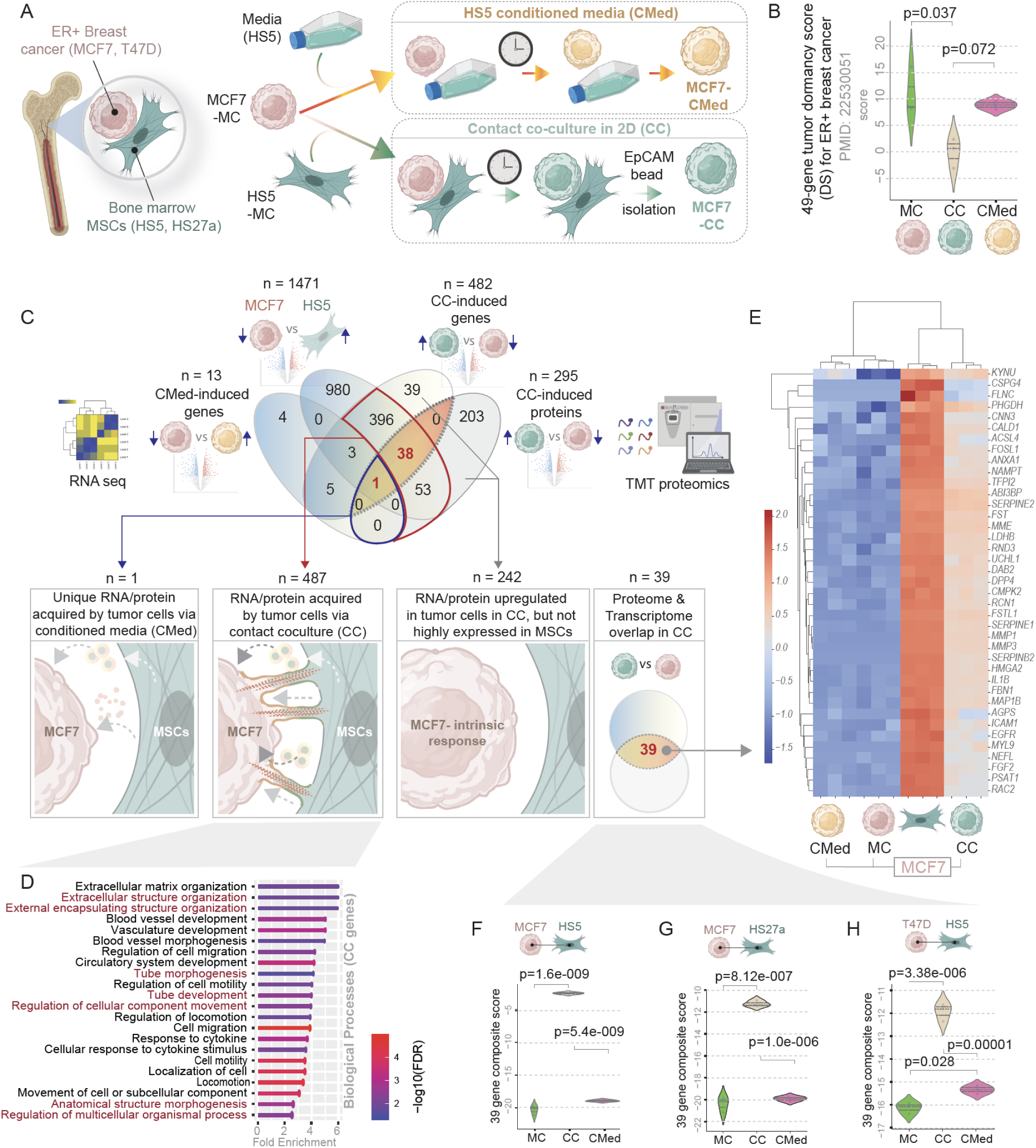
Multiomic analysis reveals RNA/protein acquired by ER+ breast cancer cells from MSCs in contact co-culture. **A.** Experimental set up for recapitulating disseminated ER+ breast cancer cells in bone marrow amidst mesenchymal stem cells (MSCs). Two different types of co-culture models were used, one with conditioned media (CMed; *top*) and one with direct contact co-culture (CC; *bottom*). MC, monoculture. **B.** Violin plots display the composite score of a 49-gene tumor dormancy score (DS) specific for ER+ breast cancers and previously validated in 4 independent cohorts to predict recurrence. **C.** *<u>Top</u>*: Venn diagram depicts multiple DEG analyses between different indicated groups that catalog suppressed genes or proteins in contact co-culture (CC) or conditioned media (CMed) and overall differential gene expression between MCF7 versus HS5 bone marrow MSCs; bottom. *<u>Bottom</u>*: Genes/proteins induced in MCF7 cells in co-culture with HS5 MSCs are binned into 3 categories (connected by arrows) based on likely mechanisms for induction. Red = 39 uniquely upregulated transcripts in cancer cells in contact co-culture also identified by proteomics. See **Supplementary Figure 1** for genes and proteins suppressed in co-culture. **D.** Gene ontology analysis (GO Biological processes) on transcripts and/or proteins acquired by MCF7 cells from MSCs during contact co-culture. See **Supplementary Figure 2** for Reactome pathway and GO enrichment analyses on genes within each category in C. **E.** Heatmap displays hierarchical unsupervised clustering of samples using the 39 genes identified in C. **F-H**. Violin plots display the composite score of the 39 genes in E in various tumor cell-MSC co-culture models, e.g., MCF7**↔**HS5 (F), MCF7**↔**HS27a (G) and T47D**↔**HS5 (H). Statistical significance for B and F-H was assessed by one-way ANOVA and the p-values are corrected for multiple comparisons using Tukey’s method.

MCF7 cells exposed to contact culture (CC), but not conditioned media (CMed), correlated with substantial lowering of a previously defined 49-gene signature for tumor cell dormancy (**Fig 1B**); low dormancy scores confer an ∼2.1-fold increase in risk of recurrence in 4 independent cohorts of patients with ER+ breast cancers [p <0.000005] (22). These findings suggest that our co-culture model captures key transcriptomic changes in ER+ breast cancer cells of translational relevance to more aggressive disease in patients.

We carried out combinatorial multiomics analyses to determine what proteins or transcripts expressed highly in MSCs are ‘borrowed’ by MCF7 cells. To this end, we first created a catalog of genes differentially expressed (DEGs; upregulated) in MSCs, but not in MCF7 cells (MCF7 vs HS5; n = 1471; **Fig 1C**). This catalog served as a denominator for all likely candidate genes that a tumor cell may lack originally but acquire from MSCs via intercellular communication. Next, we performed pairwise DEG analysis on the other MCF7 samples subjected to different co-culture conditions. Direct contact co-culture induced many genes (n = 482) and proteins (n = 295) (CC-induced genes; **Fig 1C**). Conditioned media induced few genes in cancer cells (CMed-induced genes, n = 13; **Fig 1C**). An overlay showed the following: (i) Exposure to conditioned media alone induced only a single unique protein/transcript in MCF7 cells, implying soluble mediators or exosomes as potential modes of communication; (ii) By contrast, contact culture accounted for 487 unique proteins/transcripts, implying direct contact as the major mode of tumor-MSC communication and the largest share of uniquely induced genes; (iii) Contact culture increased 242 unique proteins/transcripts (39 transcripts and 203 proteins with no proteome-transcriptome overlap) in MCF7 cells. These transcripts/proteins were not highly expressed in HS5 cells. We presumed this last category to be tumor cell-intrinsic response to contact culture, unlikely to be transferred directly from MSCs to breast cancer cells. We note that neither the experimental design nor the analytical steps can conclude definitively if transport of a given target occurred as a protein or as mRNA because unknown factors (transcript/protein half-life) likely confound such conclusions. Similarly, while we infer that 487 unique RNAs/proteins could be transferred during contact culture, i.e., “borrowed” directly by cancer cells from the MSCs, we cannot exclude some cancer cell-intrinsic contributions to the overall levels. We did observe, however, enrichment of the ‘borrowed’ RNA/proteins for biological processes that govern tube morphogenesis, organization of extracellular encapsulated structures, and movement of anatomic structures in the context of multicellular systems (**Fig 1D**). Enrichment of these processes further suggests the potential for direct transfer of RNA/protein from MSCs to cancer cells through structures such as TNTs. Reactome and gene ontology analyses of the genes induced by conditioned media, contact culture and intrinsic response present a comprehensive picture of distinct modes by which MSCs shape behavior and function of cancer cells (**Supplementary Fig 1**). A similar analysis for transcripts and proteins downregulated during contact culture and when exposed to conditioned media revealed genes/proteins that reduce the dependence on ER signaling (**Supplementary Fig 2**). We provide a full catalog of all upregulated (**Supplemental information 1**) and downregulated (**Supplemental information 2**) genes/proteins during contact culture.

### A gene signature of contact culture is prognostic and predictive in ER+ breast cancers

We noted 39 unique, upregulated genes identified by both RNA seq and TMT proteomics in MCF7 cells during contact co-cultures (**Fig 1C**). Direct contact culture uniquely upregulated almost all genes (n=38; **Fig 1E**) except for one (*KYNU*) upregulated in conditioned media. Hierarchical clustering, an unsupervised learning technique used to group similar objects into clusters, showed the 39 genes grouped the samples into two distinct groups within a dendrogram: MCF7 cells subjected to culture in direct contact with HS5 cells (CC; **Fig 1E**) clustered with the HS5 monoculture samples. By contrast, MCF7 cells cultured in conditioned media clustered with the MCF7 monoculture samples (compare MC and CMed; **Fig 1E**). A composite score of the levels of these 39 genes confirmed their statistically significant increase in contact cultures but not conditioned media (**Fig 1F**). This pattern of gene induction repeated when we swapped HS5 cells with another MSC line, HS27a, in co-cultures (**Fig 1G**) or swapped MCF7 cells with another ER+ breast cancer line, T47D (**Fig 1H**). These findings demonstrate that the 39-gene signature represents a conserved feature of how MSCs impact the properties or behavior of ER+ breast cancer.

The 39 genes showed enrichment of processes related to growth factor, PI3K/Akt, and *IL4/IL13 and IL10*-centric tolerogenic cytokine signaling (**Fig 2A**), suggesting that acquisition of these 39 genes from MSCs may impact these aspects of cancer cell biology and/or behavior, including evasion of the immune system.

**Figure 2.**
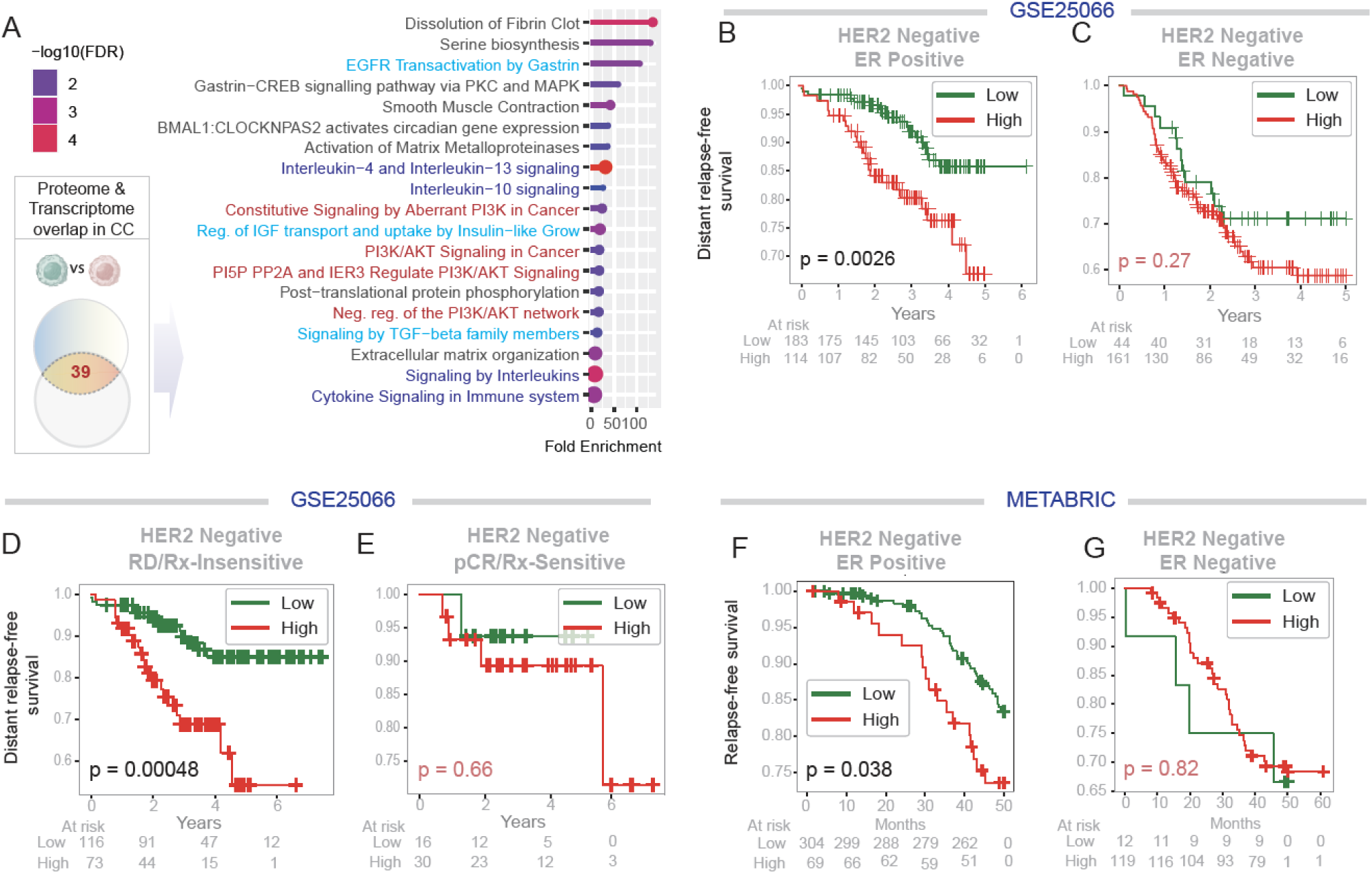
A contact culture signature derived from proteome-transcriptome overlap carries prognostic and predictive value in ER-positive breast cancer. **A.** Reactome pathways enriched in the 39 genes identified in Figure 1C. The PI3K-Akt signaling pathway (red), growth factor signaling pathways (teal blue), and the tolerogenic cytokine pathways (navy blue) are highlighted. **B-G**. Kaplan Meier survival plots on HER2 negative breast cancer patients from two independent cohorts with known outcomes over time (relapse/metastasis-free survival), stratified as high versus low expression of the 39-gene signature (see Methods). Statistical significance was assessed by Log Rank analyses. RD, residual disease; PCR, pathologic complete response; Rx, treatment.

To test translational relevance of the 39-gene signature, we applied it to a publicly available dataset that prospectively tracked outcomes of ER+, Her2-negative breast cancers following neoadjuvant taxane-anthracycline chemotherapy (23) (**Fig 2B-E**). Kaplan-Meier curves revealed that high levels of the contact culture signature correlated with significantly worse distant relapse-free survival in patients with ER+ (**Fig 2B**), but not ER-, breast cancers (**Fig 2C**). High levels of the close contact signature also predicted worse outcomes in patients with treatment resistant, but not treatment sensitive, disease (**Fig 2D-E**). We reproduced these findings in an independent Her2-negative ER+ cohort from a large dataset, i.e., METABRIC (Molecular Taxonomy of Breast Cancer International Consortium (24)), a landmark genomic and transcriptomic study of 2,000 individual breast tumors (**Fig 2F-G**). These findings establish that the 39 genes presumed to be borrowed by ER+ breast cancer cells via direct contact with MSCs carry translationally relevant information; they identify patients at greater risk for death from recurrent breast cancer.

### ‘Borrowed’ genes integrate with a cancer cell intrinsic response during contact culture

Although TMT based proteomics can reliably detect as small as ∼1/10^th^ of the changes compared to label free proteomics (25), it suffers from interference and distortion, which disproportionately impact low abundance proteins that often are the ones borrowed (26, 27). To circumvent this limitation and improve detection of relevant ‘hits’, we utilized a protein-protein interaction (PPI) network approach to identify relevant proteins, based on the assumption that the proteins function through meaningful interactions with each other. Briefly, we used the 39 proteins as ‘seeds’ to build a PPI network comprised of 2743 proteins (**Fig 3A**). We overlayed the network-derived list with ‘borrowed’ (n=487) and ‘intrinsic’ (n=242) gene or protein sets (see **Fig 1B**) acquired by tumor cells via contact culture (n=487) and genes or protein sets upregulated in tumor cells in contact culture but not highly expressed in MSCs. Generating the proteome network using the STRING database (see *Methods*) and then intersecting the proteome with the differential transcriptome increased signals (i.e., ‘hits’) by leveraging the sensitivity of transcriptomics and specificity of proteomics. This process produced a more focused network using 159 genes that ER+ breast cancer cells borrow from MSCs and 76 genes in the cancer cell intrinsic response group. **Fig 3C** displays the multilayer network connecting borrowed (pink) and intrinsic response (green) genes. Within the PPI network, 1482 nodes/proteins interacted within the ‘borrowed’ layer and 835 nodes/proteins interacted within the intrinsic layer. The borrowed response interactome included Akt, growth factors and/or their receptors (*EGFR, VEGFA, WNT5A*), and integrins (**Fig 3C**), as noted earlier (**Fig 2A**). The ‘borrowed’ network also revealed meaningful enrichment of other candidates, including the gap junction protein, *GJA1*/CX43, which has been implicated in establishing TNTs that facilitate exchange of molecules (28). The two layers shared 567 common nodes/proteins, indicating meaningful connectivity between two layers (*p* value of 0.00096 by hypergeometric test). The shared nodes were enriched for three prominent themes: membrane trafficking, immune signaling, and growth factor signaling (**Fig 3D**).

**Figure 3.**
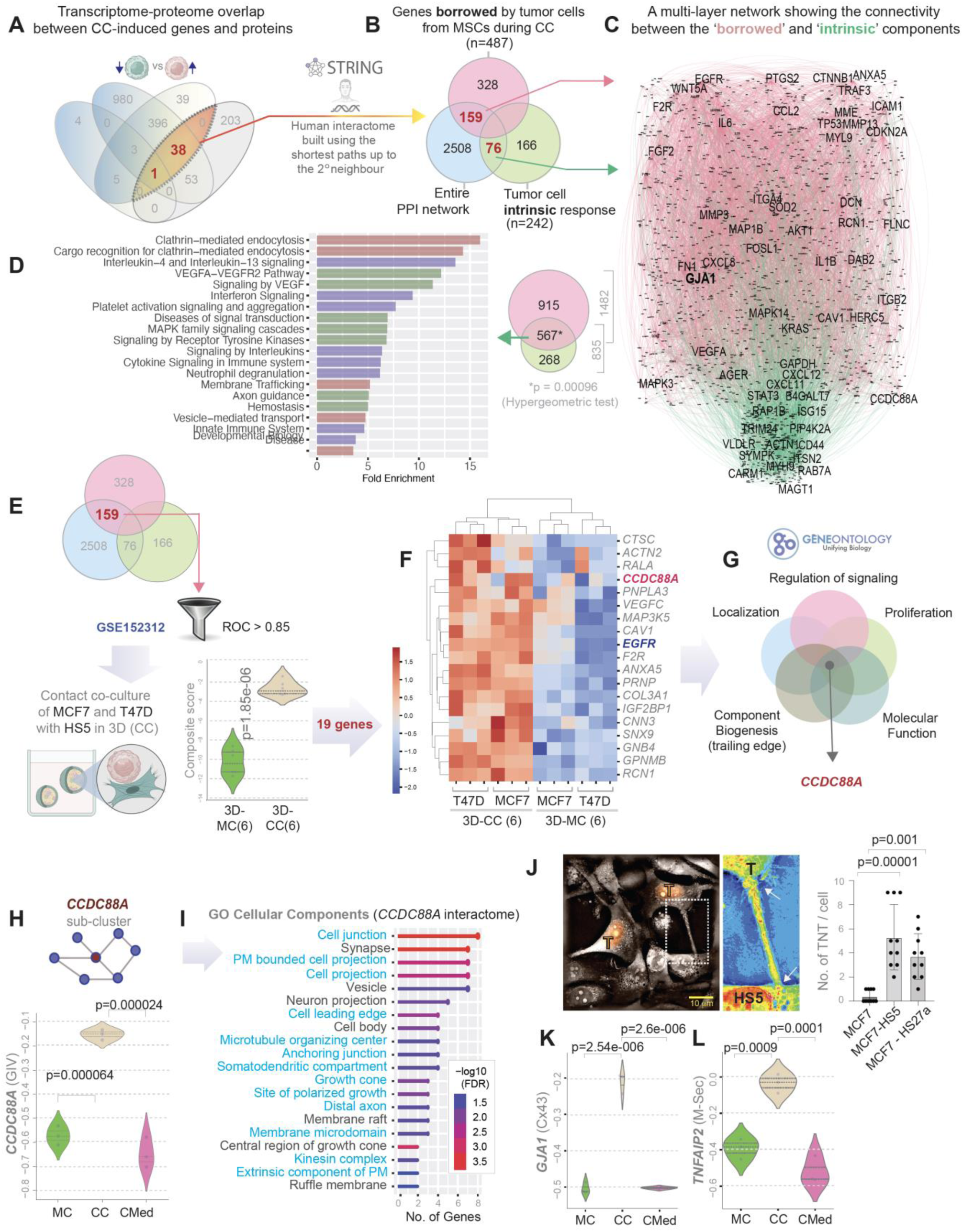
Network analysis pinpoints GIV as a central orchestrator of the co-culture borrowed gene signature. **A.** Workflow for protein-protein interaction (PP network analysis using the 39 gene signature from Figure 1E as ‘seeds’ to build a PPI network using the STRING database. **B.** Venn diagram shows overlaps between the PPI network and the candidate genes/proteins identified in Figure 1C as either likely to be ‘borrowed’ from MSCs or upregulated as part of cancer cell’s ‘intrinsic’ response during contact co-cultures. **C.** A multi-layer PPI network shows connectivity between ‘borrowed’ (n = 159) and ‘intrinsic’ (n = 76) proteins with key nodes in the network labeled. **D.** Reactome pathways (*left*) enriched in the 567 proteins shared between the borrowed and intrinsic layers of the PPI network in C. Venn diagram (*right*) shows the total nodes in each layer and degree of overlap. **E.** The 158 ‘borrowed’ genes identified in our 2D co-culture models were further filtered using a public dataset from 3D co-culture models with MCF7 and T47D cells grown with HS5 MSCs. A threshold ROC AUC > 0.85 identified 19 genes. A violin plot shows the composite score for expression of 19 genes in the 3D mono (MC) vs contact (CC) cultures. **F.** Heatmap of z-score normalized expression of the 19 genes. MC, monoculture; CC, contact culture. **G.** Venn diagram summarizes gene ontology (GO) analyses, identifying *CCDC88A* as an invariant player across diverse biological processes enriched within the 19 genes. **H-I**. Violin plot (H) shows induction of *CCDC88A* in MCF7 cells when subjected to contact coculture (CC) but not conditioned media (CMed). Statistical significance was assessed by Welch’s t-test. Only significant p values are displayed. Graph (I) displays the GO cellular components enriched within the *CCDC88A*/GIV-subcluster in the PPI network in C. Components highlighted in blue indicate processes required for long-range intercellular transport via tunneling nanotubes. **J**. Images (left) obtained by interferometry microscopy of MCF7 cells (red nuclei; “T”) in contact coculture with MSCs (HS5) with nanotube connecting cells. The boxed area is magnified on the right. Arrows mark ends of the nanotube. **Supplementary Figure 3** shows a montage of images of MCF7 cells in mono-(MC) or contact coculture (CC) with HS5 MSCs. Bar plots (right) show quantification of the average number of TNTs observed in each condition per high power field (HPF). **K-L**. Violin plots show induction of *GJA1* (K) and *TNFAIP2* (L) in MCF7 cells when subjected to contact co-culture (CC) but not in conditioned media (CMed). Statistical significance for E, H, J-L was assessed by one-way ANOVA and the p-values are corrected for multiple comparisons by using Tukey’s method.

Findings suggest functional integration between the two components; the induction of one (the intrinsic cancer genes) may either enable functions of and/or reflect consequences of the other (the borrowed genes). **Supplemental Information 3** presents the edge and node list of the entire PPI network and various subnetworks.

### *CCDC88A*/GIV as a key gene borrowed by cancer cells

To validate and/or characterize key components of the borrowed transcriptome/proteome and yet reduce the risk of noise (of STRING database) and artifacts (of 2D culture), we further refined this list (n = 159) generated using 2D cultures by leveraging a published dataset using the same cell lines as us except in 3D cultures (GSE152312) (**Fig 3E**) (29). We carried out this refinement with a two-fold intention: (i) to enhance the physiological context and translational relevance because TNTs can form in 3D culture systems (30); and (ii) to provide continuity between 2D and 3D TNT biology, which has been difficult to achieve in a field still in its infancy (31). Refinement gave a handful of genes (n=19) also induced during 3D close contact culture in both MCF7 and T47D cells (**Fig 3F**). We noted that two of the 19 genes (*CCDC88A* and *EGFR*) are key components of a recently described phenomena, growth signaling autonomy, which endows breast tumor cells with plasticity and stemness (among other features) especially during hematogenous dissemination in metastasis (32, 33). Gene ontology (GO) analysis of these 19 genes revealed that *CCDC88A* [which encodes the protein Gα-Interacting, Vesicle-associated (GIV or Girdin)] is commonly shared among key processes upregulated in ER+ breast cells during reprogramming of signaling to adopt a mesenchymal state (34–38) (**Fig 3G**). Compared to MCF7 monocultures, 2D direct contact cultures, but not conditioned medium, markedly upregulated *CCDC88A* (**Fig 3H**). Focused analysis of the *CCDC88A/GIV*-subnetwork from the multi-layer interactome (**Fig 3C**) revealed that the interactome of GIV could support numerous pathways and processes involved in cell projections and membrane domains (**Fig 3I**), suggesting that *CCDC88A* associates with direct intercellular contacts.

TNTs constitute a major pathway to transfer materials, including RNA and proteins, between cells of the same and different types(39) through direct intercellular contacts. Using interferometry microscopy, we identified numerous TNTs connecting MCF7 and HS5 cells (**Fig 3J**, **Supplementary Figure 3**). Images revealed significantly more heterotypic TNTs connecting MCF7 cells with HS5 or HS27a stromal cells than homotypic TNTs connecting MCF7 cells in monocultures (**Fig 3J**-*right*). TNT formation in contact cultures also associated with elevated expression of two proteins previously implicated as central factors essential for their formation: (i) the gap junction protein *GJA1* (28) (connexin 43, CX43; **Fig 3K**) and the TNT marker, *TNFAIP2* (40) (M-Sec; **Fig 3L**). To investigate if gap-junction or TNT-facilitated intercellular communication is functional, we conducted the well-established calcein-AM transfer assay (41, 42). Co-culture of MCF7 cells stably expressing mCherry (recipients) with calcein-labeled HS5 MSCs (donors) or calcein-labeled MCF7 cells showed successful transfer of calcein into mCherry-labeled MCF7 cells (**Supplementary Figure 4A**). Both donor cells (HS5 and MCF7 cells) transferred calcein to recipient MCF7 cells. Although MCF7 cells formed both heterotypic and homotypic TNTs, transfer is markedly higher from MSCs (**Supplementary Figure 4A**). We also detected transfer from MCF7 cells to HSCs, albeit at lower efficiency (**Supplementary Figure 4B**). These results confirm that our 2D contact cultures allowed intercellular transport.

Together these findings suggest that TNTs in contact cultures could provide a route for breast cancer cells to borrow from MSCs certain RNAs and proteins, among which *CCDC88A*/GIV may be a central player in functional outcomes.

### Transport of GIV(*CCDC88A*) from MSCs to ER+ breast cancer cells

Although prioritized through computational rationale (**Fig 3E-I**), we selected GIV (**Fig 3F**) because of three key reasons: *<u>First</u>*, as a large multi-modular protein that integrates signals from diverse classes of receptors and signaling pathways, GIV drives tumor cell aggressiveness (stemness (43), survival after DNA damage (44), invasiveness (44), chemoresistance (45), and acquisition of metastatic potential (46)) in multiple cancers, including breast (47), and is considered to be a metastasis-related protein (48) (**Fig 4A**). *<u>Second</u>*, unlike stromal cells (49, 50) and triple negative breast cancers, which express high levels of GIV, ER+ breast cancer cells, such as MCF7 and T47D, lack GIV (51) due to an alternative splicing event [intron 19 retention] that leads to the loss of the resultant transcript to non-sense mediated decay. The concept of haves (MSCs) lending GIV to have nots (ER+ tumor cells) for potential gain in metastatic potential by the latter appeared interesting. Consistent with this concept, prior studies demonstrated that patients with ER+ breast cancers do express GIV, and such expression correlates with more metastasis and poorer survival (52, 53). *<u>Third</u>*, GIV has recently been shown to be a key orchestrator of secretion-coupled autonomy (32), wherein cancer cells may secrete and sense their own growth factors to survive. Proteomic studies confirmed that GIV enabled autocrine growth factor signaling autonomy specifically within the EGF/EGFR system(32). We note that EGFR is one of the initially identified list of 39 ‘borrowed’ RNAs and proteins in contact culture (**Fig 1E**). EGFR remained on the list even after the 3D contact culture refinement (**Fig 3E**) as one of the 19 “borrowed” genes, alongside *CCDC88A* (**Fig 3F**).

**Figure 4.**
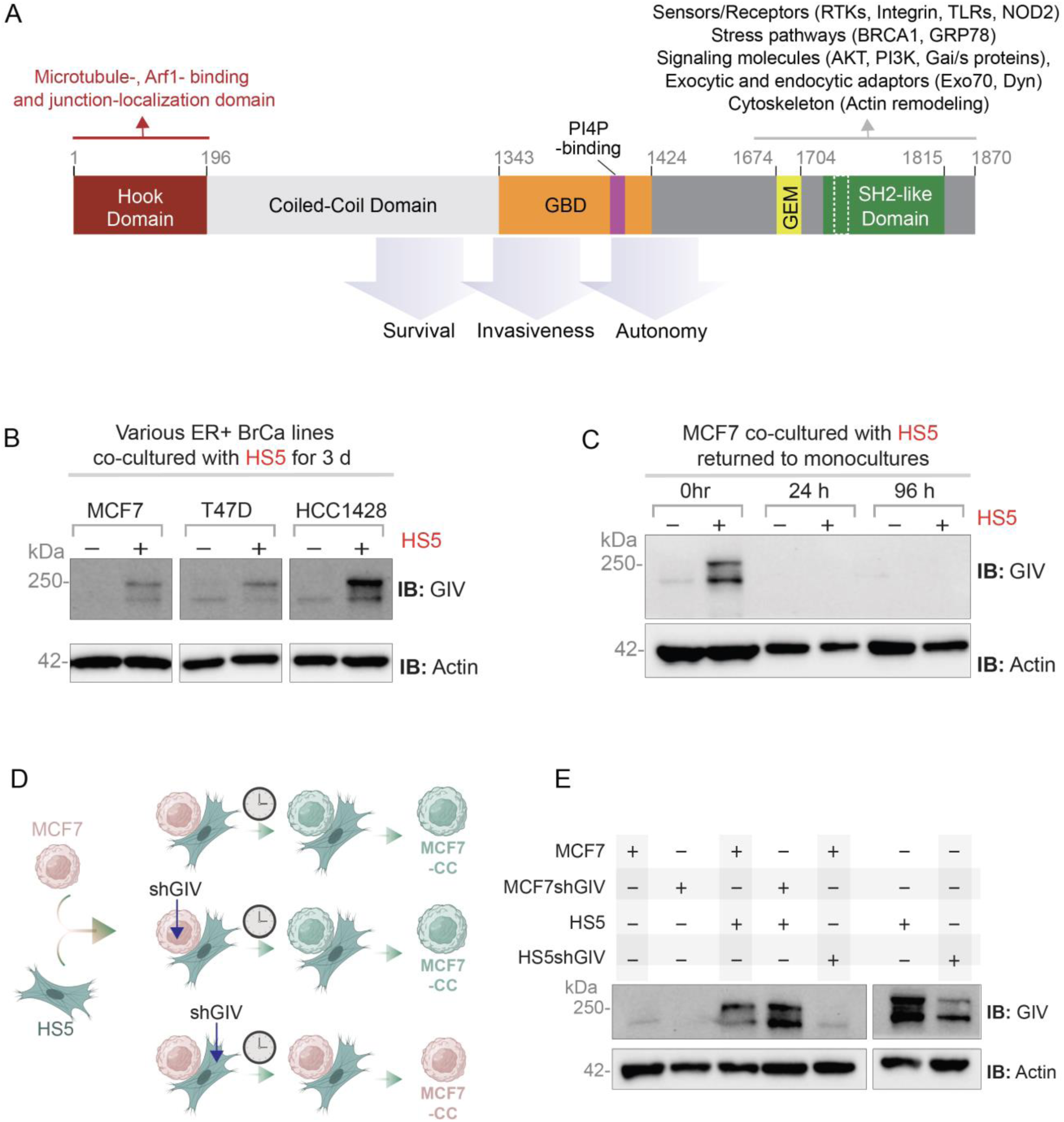
GIV transport from MSCs to ER+ breast cancer cells requires ongoing interactions. **A**. Schematic illustrates key functional domains of GIV. The C-terminus contains multiple short linear motifs that enable diverse tumor-promoting interactions (receptors, signaling molecules within diverse pathways, and components of membrane trafficking). GBD, G protein binding domain; GEM, guanine nucleotide exchange modulator; SH2, Src-like homology domain. **B**. Equal aliquots of lysates of 3 ER+ breast cancer cells (MCF7, T47D or HCC1428) in monoculture or recovered after co-culture with HS5 cells were analyzed for GIV and actin (as a loading control) by immunoblotting (IB). **C**. Equal aliquots of lysates of MCF7 cells prepared either immediately after recovery from co-cultures with HS5 cells (0 h) or after an additional 24 or 96 h of culture after removal from HS5 contact were analyzed for GIV and actin (as a loading control) by IB. **D-E**. Schematic (D) outlines the key manipulations (i.e., treatment with shRNA for GIV) done to either MCF7 or HS5 cells in contact co-cultures. Equal aliquots of MCF7 cells recovered from the co-cultures (left) or HS5 cells in monocultures were analyzed for GIV and actin (loading control) by IB.

GIV protein was virtually undetectable in lysates from monocultures of 3 different ER+ breast cancer cells but detected as a full-length protein (∼250 kDa) in cells recovered with EpCAM-based immunocapture and lysis following 3 days in contact cultures with HS5 cells (**Fig 4B**). GIV protein was detected by immunoblotting in MCF7 cells lysed immediately after separation from HS5 cells but not after 24 to 96 hours of additional culture post separation (**Fig 4C**). These findings indicate that once acquired, cancer cells rapidly lose GIV after return to monoculture. ER+ breast cancer cells require ongoing close contact with MSCs to maintain levels of GIV.

To confirm if MCF7 cells acquire GIV from HS5 cells and determine if transport occurred as transcripts or proteins, we stably expressed short hairpin (sh) RNA targeting GIV in either MCF7 or HS5 cells and put them in contact culture with each other in various combinations (see **Fig 4D**). Batch populations of cells showed ∼80% depletion of GIV in HS5 cells (**Fig 4E-***right*). Contact cultures of control MCF7 and HS5 cells showed expected upregulation of full-length GIV in MCF7 cells (**Fig 4E-** *left*; lane 3). Contact cultures of MCF7-shGIV and HS5 cells still showed increased amounts of GIV protein, indicating that shRNA against *CCDC88A* did not reduce GIV transfer (**Fig 4E-***left*; lane 4). However, MCF7 cells co-cultured with HS5-shGIV cells did not demonstrate GIV protein transfer (**Fig 4E-***left*; lane 5). Because MCF7 cells expressing shGIV continued to express GIV protein, findings indicate that the observed upregulation of GIV in MCF7 cells in contact culture with HS5 cells relies predominantly on transfer of GIV protein. Some transfer of GIV as mRNA cannot be ruled out.

### Cx43 and GIV interact and are co-transported through intercellular communication sites

Because gap junctions serve as sites for TNT formation and/or attachment (30, 54) and CX43 is a shared molecular entity for both, we assessed CX43 expression in MCF7 cells that were either in monocultures or recovered from contact cultures with HS5 cells. Co-culture with HS5 cells increased CX43 in MCF7 cells, while monocultures of MCF7 cells showed undetectable levels of this protein (**Fig 5A**). Next, we used two perturbagens to disrupt intercellular communication via gap junctions: (i) carbenoxolone, a widely used chemical inhibitor of GJIC; and (ii) fulvestrant, which inhibits transcription of ER targets, including CX43 (55, 56). Both perturbagens virtually eliminated upregulation of GIV and CX43 in MCF7 cells co-cultured with HS5 cells (**Fig 5B**), while these compounds had minimal effects on CX43 in HS5 cells alone except at 50 µM carbenoxolone (**Supplementary Figure 5**). These results show that MCF7 cells require direct, CX43-mediated intercellular communication to borrow GIV protein from HS5 cells.

**Figure 5.**
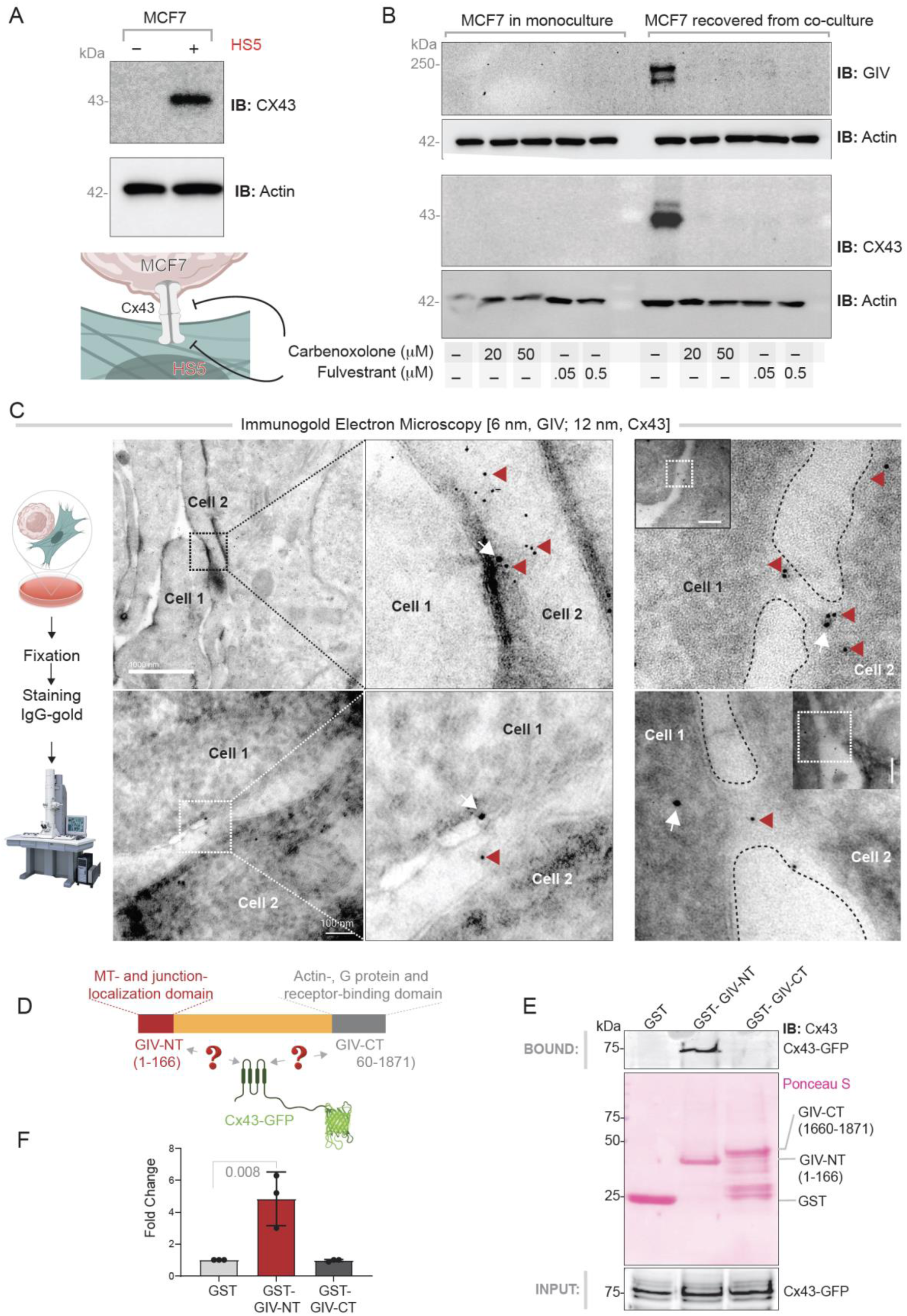
Cx43 and GIV interact and are co-transported through intercellular communication sites. **A**. Equal aliquots of lysates of MCF7 cells in monoculture or recovered after co-culture with HS5 cells were analyzed for Cx43 and actin (as a loading control) by immunoblotting (IB). **B**. MCF7 cells in monoculture or in co-culture with HS5 cells were treated with the indicated concentrations of carbenoxolone or fulvestrant prior to lysis. Equal aliquots of lysates were analyzed for GIV, Cx43 or actin (the latter as a loading control) by IB. See **Supplementary Figure 5** for similar studies on HS5 monocultures. **C**. Electron micrographs of immunogold stained MCF7↔HS5 contact co-cultures. Red arrowhead denotes 6 nm gold particles (GIV); White arrow denotes 12 nm gold particles (Cx43). Scale bar = 100 µm. **D-F**. Schematic of constructs (D) used in pull-down assays with recombinant GST-tagged GIV N- or C-term fragments or GST alone (negative control) proteins immobilized on glutathione beads and lysates of Cos7 cells transiently expressing GFP-Cx43. Bound Cx43 was visualized by immunoblotting (E-top). Equal loading of GST proteins was confirmed by Ponceau S staining (C-bottom). Bar graphs (F) display the relative binding of Cx43 to various GST proteins. Data are represented as mean ± SEM (n = 3); p-value determined by one-way ANOVA.

To resolve the nature of these CX43-dependent compartments that facilitate intercellular communication, we carried out ultrastructural analysis by immunogold electron microscopy. CX43 and GIV co-localized near gap junctions (**Fig 5C***-top left*), at/near more elaborate and less defined contact sites (**Fig 5C***-bottom left*), or within tube like passages connecting cells (**Fig 5C***-right*).

Co-localization of CX43 and GIV suggested that these proteins interact. Using CX43-GFP expressed in Cos7 cells and purified GST fusions of the N-terminal or C-terminal part of GIV in pull down assays, we confirmed that the N-terminus, but not the C-terminus of GIV binds directly to CX43 (**Fig 5D-F**). This finding is consistent with prior work demonstrating the importance of GIV-NT in junctional localization (57).

Together, these findings suggest that CX43 and GIV interact and co-migrate from MSCs to ER+ breast cancer cells during CX43-mediated intercellular transport.

### Adding GIV to ER+ breast cancer cells recapitulates key functions gained during contact culture

We asked to what extent GIV alone accounts for or recapitulates functions gained by ER+ breast cancer cells during contact culture with MSCs. We generated MCF7 cells stably expressing GIV and confirmed that exogenous expression produced levels of GIV comparable to that from intercellular transfer from HS5 cells in contact co-cultures (**Fig 6A**) (**Supplementary Fig 6A**). RNA sequencing of MCF7-GIV cells revealed that ∼20% of the genes in the ‘borrowed’ (n = 32 genes) and ‘intrinsic’ response (n = 17 genes) components of contact culture perfectly classified MCF7 cells with or without GIV (**Fig 6B, C**). We present a complete catalog of these genes in **Supplemental Information 4**. Heat maps display the differential expression of these genes (**Fig 6E, F**). Of the tumor intrinsic gene response, we previously discovered that co-culture with MSCs upregulated *LCN2,* an iron sequestering protein, and *CD44,* a marker of stem-like cells, in breast cancer cells (19). Pathway analysis of the 32-gene borrowed signature recapitulated by stable expression of GIV alone showed enrichment for numerous processes related to response to external stimuli, while the 17-gene signature included vesicular transport and exocytosis (**Supplementary Fig 6B-C**). These data reinforce our focus on GIV within the list of genes transferred from MSCs to breast cancer cells in bone marrow to enhance aggressive traits and facilitate disease progression.

**Figure 6.**
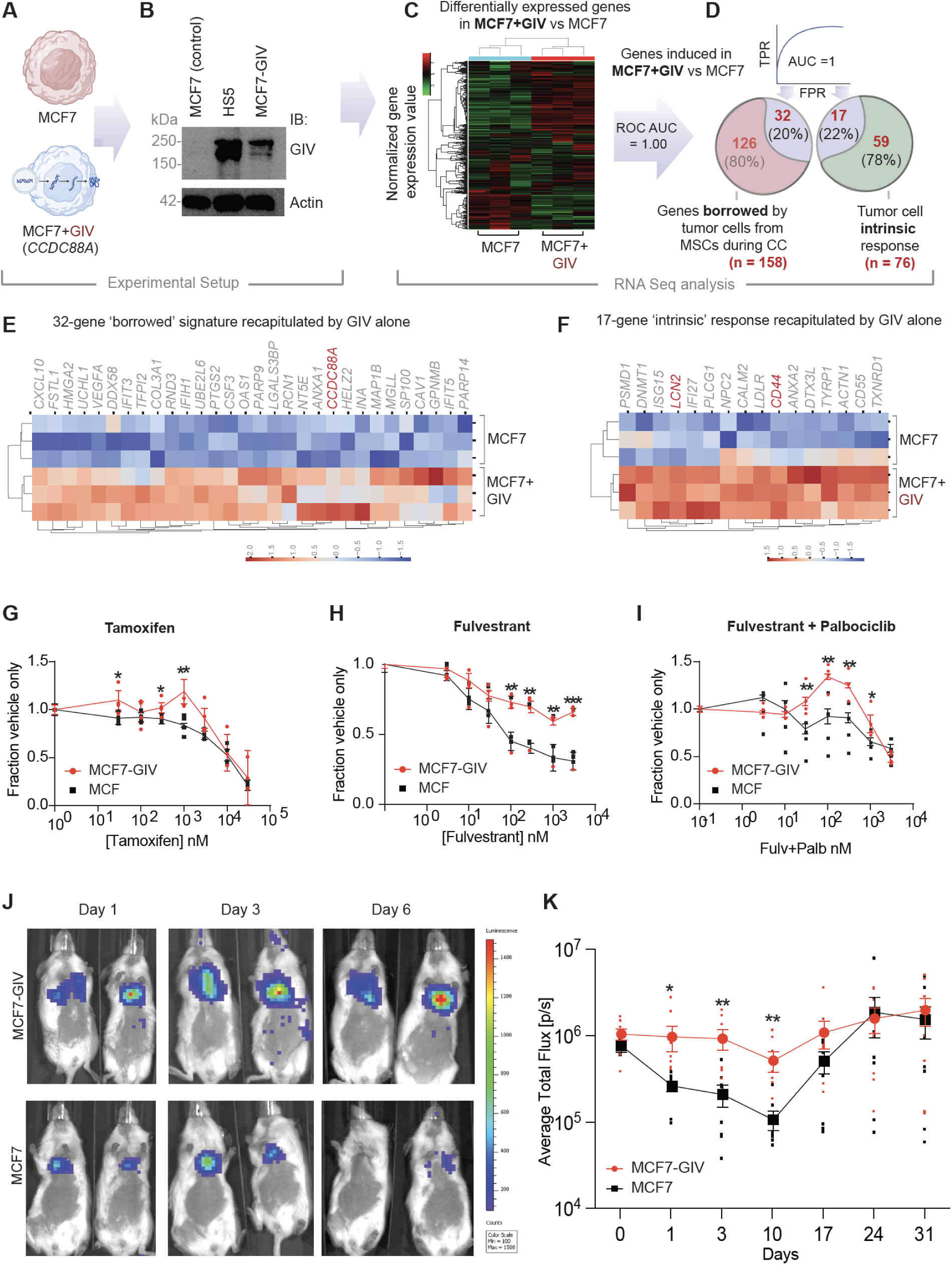
Expression of GIV recapitulates the MSC close contact signature; confers resistance to ER-targeted therapies; and promotes early dissemination of ER+ breast cancer cells. **A**. Schematic shows creation of a stable MCF7 cell population expressing exogenous GIV (*CCDC88A*) by a piggyBac transposase vector. **B**. Immunoblots of whole cell lysates of cells in A, confirming expression of full-length GIV. **C-D**. Workflow (C) of RNA seq and normalized gene expression analysis of cells in B analyzed for the ‘borrowed’ (n=158) and ‘intrinsic’ (n=76) genes to perfectly classify MCF7-GIV cells from control MCF7 (ROC-AUC 1.00). Venn diagram (D) shows genes induced with GIV alone as % of the total ‘borrowed’ and ‘intrinsic’ signatures. **E-F**: Heatmaps display hierarchical unsupervised clustering of MCF7 and MCF7-GIV cells by z-score normalized gene expression for the subset of genes induced among the borrowed (E) and intrinsic (F) signatures. **Supplementary Figure 6** shows reactome pathway analysis of these genes. **G-I**. Graphs display viability for MCF7-GIV and control MCF7 cells exposed to increasing concentrations of tamoxifen (G), fulvestrant (H), or fulvestrant and 100 nM palpociclib (I). Data are displayed after normalization to cells treated with vehicle only. **J-K**. Equal number (1 x 10^5^) of MCF7-GIV or control MCF7 cells were injected into the left cardiac ventricle of 7-10-wk-old female NSG mice (n=8 per group). Representative bioluminescence images (J) show mice at days 1-, 3-, and 6 post injection. G-I and K Graphs show mean values ± SEM of each group at specific data points. Statistical significance between two groups at each time point was computed using repeated measures ANOVA.*, p < 0.1, **, p <= 0.05 and, ***, p<=0.01 indicates corrected p-values using Tukey’s method.

To investigate effects of GIV on drug resistance, we focused on anti-estrogen drugs tamoxifen (a selective estrogen receptor modulator) (58) and fulvestrant (a selective estrogen receptor degrader) (59). We also tested combination therapy with fulvestrant and palbociclib, an inhibitor of CDK4/6 used with anti-estrogen therapy for patients with metastatic ER+ breast cancer. Following three days of treatment in low serum, physiologic glucose medium, we quantified relative numbers of viable cells at various concentrations of tamoxifen, fulvestrant, or fulvestrant plus a fixed concentration of palbociclib. MCF7-GIV cells showed resistance to each of these treatment regimens, particularly fulvestrant (**Fig 6G-I**). These data establish that GIV confers resistance to clinically used drugs for ER+ breast cancer and suggest that upregulation of GIV in cancer cells in close contact with MSCs may permit disseminated ER+ breast cancer cells to survive therapy in bone marrow.

Because GIV coordinates intracellular signaling pathways enabling growth-factor autonomous survival and proliferation of circulating tumor cells (32, 60), we investigated effects of GIV on hematogenous dissemination of ER+ breast cancer cells. We injected MCF7 and MCF7-GIV cells into the left cardiac ventricle of female immunodeficient NOD.Cg-Prkdc scid (NSG) mice and tracked cancer cells using bioluminescence (**Fig 6J**). Viable MCF7 cells dropped steadily from day 1 through day 10, while no significant decrease occurred in mice injected with MCF7-GIV cells. From days 0-1, signal from control MCF7 cells decreased by approximately 70%, while MCF7-GIV cells diminished by only approximately 10%. From days 1-3, losses of viable cells were approximately 20% and <10% for MCF7 and MCF7-GIV cells, respectively. The rapid drop in MCF7 cells is not unexpected because we did not supplement mice with estrogen, which typically is required for MCF7 cells to proliferate in vivo. Intriguingly, we noted that differences between groups diminished over time from days 10-24 and became virtually indistinguishable thereafter. These results reveal that GIV facilitates survival during the early stages in dissemination of ER+ breast cancer cells. Because contact culture studies in vitro take ∼3 days to transfer GIV, it is possible that MCF7 cells borrowed GIV from stromal cells that have abundant amounts of the protein.

### Tumor cells may acquire *CCDC88A* and other ‘borrowed’ genes also from fibroblasts

Studies have documented that disseminated tumor cells and metastases in one organ can seed metastases in a second site, emphasizing the need to understand how the bone marrow environment promotes aggressive phenotypes in breast cancer (61). We asked if the catalog of ‘borrowed’ genes we describe here maintained relevance beyond MSCs and extended also to MSCs that can be recruited to the tumors where they give rise to cancer-associated fibroblasts (CAFs (62–64)). CAFs establish TNTs with cancer cells (65), express high levels of GIV, and rely upon GIV to aid cancer progression (66). We leveraged the only publicly available, published study that compared monocultures of MCF7 cells with contact cultures of MCF7 cells and BJ fibroblasts by RNA seq (67). As a third comparator group, this prior study cultured MCF7 cells in extracellular matrix (ECM) derived from conditioned media from the BJ fibroblasts (CMed) (**Fig 7A**). Contact culture, but not ECM-treated growth conditions, induced each of the 3 borrowed gene signatures we described earlier (**Fig 7B-F**). Most importantly, *CCDC88A* was among the “borrowed” gene signature induced during contact culture with fibroblasts. Furthermore, we found that xenografted tumor cells depleted of GIV by CRISPR Cas9 (**Supplementary Figure 7A**-B) gained GIV protein (**Supplementary Figure 7C**), but not mRNA (**Supplementary Figure 7D**), presumably from high-GIV expressing mouse stromal cells and/or co-implanted human mammary fibroblasts (arrows, **Supplementary Figure 7C**). These results suggest that ER+ tumor cells may acquire GIV via intercellular transport from diverse stromal cells that express GIV at high levels. This raises the possibility that the phenomenon of GIV acquisition via TNTs may not be limited to bone marrow but extend also to the primary and metastatic sites, where prolonged contact and transfer could occur between tumor cells and CAFs. It also provides the basis for the observed prognostic significance of the 39-gene signature in ER+ primary tumors (**Fig 2B, D-E**).

**Figure 7.**
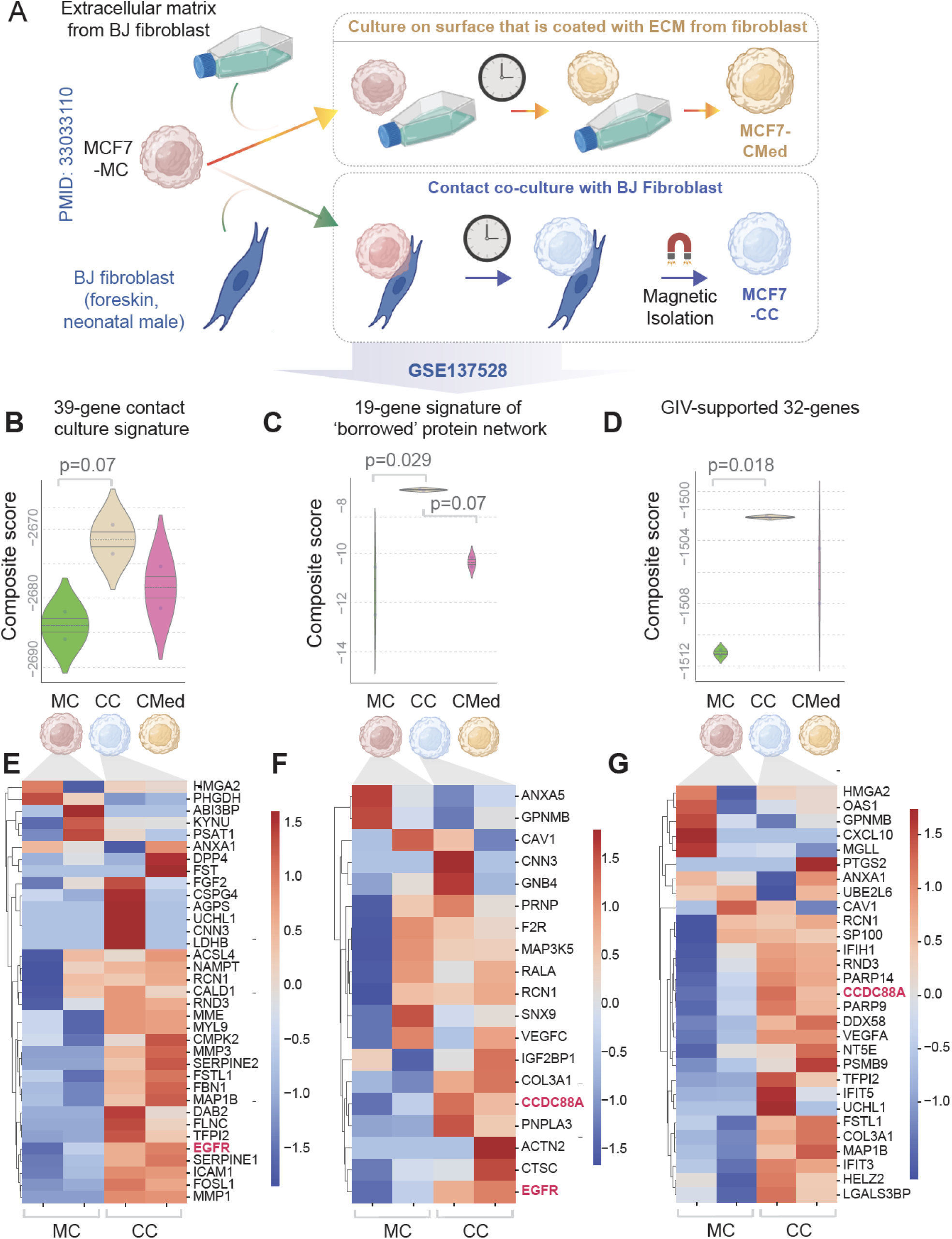
MCF7 cells borrow *CCDC88A* and other key genes during contact culture with fibroblasts. **A.** Schematic of the experimental setup in which MCF7 cells were grown either as monoculture (MC), or in direct contact cultures with BJ fibroblasts (CC) or cultured on ECM produced by the BJ fibroblasts (CMed). **B-D**. Violin plots show the degrees of induction of 39-gene 2D contact culture signature (**B**), the 19-gene signature of network of ‘borrowed’ proteins/transcripts that survived refinement through 3D contact cultures (**C**), and GIV-supported ‘borrowed’ transcriptional program of 32 genes (as in Figure 6D). Statistical significance was assessed by one-way ANOVA and the p-values are corrected for multiple comparisons by using Tukey’s method. **E-G**. Heatmaps show the induction pattern of each gene within the signatures described in B-D. Color key denotes z-score normalized counts per million (cpm).

## Discussion

The major problem we address here is the widely recognized, but poorly understood, means by which bone marrow harbors disseminated tumor cells in ER+ breast cancer, permitting long-term survival of these cells despite hormone targeted therapy (68). Persistence of disseminated tumor cells poses a documented, progressively increasing risk of delayed recurrence that most often occurs in bone (69–71) (∼50%) for patients with ER+ breast cancer. MSCs have been implicated as key enablers of tumor cells in the bone marrow niche (61, 72). However, mechanisms that support disseminated tumor cells in bone marrow and potentiate potential recurrence remain unclear. The compendium of TNT tumor-stromal transport we reveal here provides a wealth of potentially actionable targets to disrupt MSC niches for cancer cells and end the current impasse in treating bone metastases. There are potentially three major ways in which this work advances the field of cancer biology.

### A catalog of MCS-to-tumor gene transfer

First, we presented an integrated omics workflow that identifies meaningful ‘hits’ by optimally leveraging the sensitivity (transcriptomics) and specificity (proteomics) of approaches and maintains the relevance and context (2D and 3D datasets). The workflow generated a comprehensive catalog of genes transferred from MSCs to ER+ breast cancer cells during direct contact culture. The approach ensured that low levels of borrowed proteins (such as Cx43 and GIV) were not overlooked due to limitations of proteomics, whereas the 2D versus 3D comparisons ensured that the catalog is likely to be physiologically relevant and reproducible. The multiomic rigor we employed went beyond our own datasets generated using all 4 possible combinations of two ER+ breast cancer cell lines and two MSC lines. This catalog not only provides a prioritized list of genes for the scientific community to pursue but also provides knowledge of molecular mechanisms that reprogram breast cancer cells in bone marrow. For example, while MSCs regulate breast cancer cells through soluble mediators such as cytokines (73), work done by others and us revealed that key functional changes in ER+ breast cancer cells required direct close contact with MSCs (72). The present catalog provides a multiomic resource to begin to dissect how MSCs shape disease phenotypes in ER+ breast cancer and to what extent effects rely upon intercellular transfer of key molecules (‘borrowed’). We show that the borrowed components integrate seamlessly with a second set of gene expression changes that reflect the cancer cell ‘intrinsic’ responses; together, they establish a robust connectome. Key borrowed components belong to the growth factor and pro-survival PI3K-Akt signaling pathways, while the key intrinsic response is augmentation of VEGF signaling, which supports proliferation and angiogenesis. The key process suppressed is dependance on estrogen signaling, consistent with environmentally mediated resistance to hormone targeted therapies (74). The ‘borrowed’ gene signature showed demonstrable translational relevance in that it prognosticated and predicted outcome in patients with ER+ breast cancers, especially those with treatment-resistant disease. We conclude that the catalog of genes may represent high value targets for preventing MSCs from empowering tumor cells that lie in the bone marrow niche. Contact co-culture of ER+ breast cancer cells and fibroblasts reproducibly upregulated several key genes from this list, implying that findings may have broader relevance in the context of cancer cells in other tissues. Resistance to therapies for ER+ breast cancer and enhanced survival of disseminated MCF7 cells stably expressing GIV recapitulate key features of MCF7 cells in co-culture with MSCs that we reported previously (19, 21), and future studies will delineate if and how GIV alone may impact additional reported phenotypes, i.e., metabolic plasticity (20).

### Gene transfer occurs through a nanotube-based connectivity grid

Second, the catalog of genes provides valuable clues into the predominant mode of intercellular communication between MSCs and tumor cells. Intercellular protein transfer encompasses several different biologic processes and cellular structures that establish dynamic communication networks among cells in direct contact or proximity: Gap junctions, TNTs, extracellular vesicles, trogocytosis, and direct cell fusion are among such processes (75). Gene ontology analyses of our catalog of borrowed genes hinted at formation of extracellular tube-like structures, such as TNTs, as a likely mode of transport. This is consistent with prior reports of MSCs establishing TNTs with multiple types of cancer cells, including MCF7 and other breast cancer cell lines (76). Discovered originally in 2004 (14) and critiqued initially by some as artifacts of 2D culture (77), TNTs are now increasingly recognized as conduits for cell-cell communication in both physiology and disease (28, 30, 54, 78). TNTs are F-actin rich structures with junctional proteins at one end (79, 80); TNTs couple with adjacent cells through gap junctions (81–84), and both share CX43 as common molecular component. We observed formation of abundant TNTs connecting MCF7 cells with MSCs and demonstrated that they are functional using pharmacologic inhibitors, implicating gap junctions and CX43 as facilitators of transport in our cultures. The presence of EGFR on the list of ‘borrowed’ genes is intriguing. While we did not investigate transfer of this receptor as RNA or protein, transmembrane and membrane bound proteins have previously been reported to be transferred via TNTs (85). Most of the borrowed genes, however, represent cytoplasmic proteins and/or organelle- or cytoskeleton-associated proteins, which are known to use TNTs as a route for intercellular protein transfer (30, 54, 78). That the 483 borrowed genes carried an overwhelming message of intercellular transport structures (tube morphogenesis and development) suggests selection of these genes by their ability to facilitate or support TNT formation, stabilization, and transport. Integration of the borrowed and intrinsic genes through PPI networks indicates that the nature of the exchange is meaningful and advantageous to cancer cells (and potentially also to MSCs). The TNT-dependent tumor-MSC communication we reveal here resembles nanotubes (86, 87) and pili-dependent communication in pathogenic biofilm-producing microbes (which are responsible for ∼80% of chronic infectious diseases) (88). Unicellular pathogens deploy these structures to communicate with each other, diverse host cells, or other pathogens to ultimately produce an extracellular grid to adopt a ‘multicellular lifestyle ‘(89, 90). Such adaptation promotes drug resistance, survival, transfer of resistance genes, and metabolic dormancy, in effect, making bacterial biofilms more difficult to eradicate than when present as unicellular organisms (reviewed in) (91, 92).

### Acquisition of GIV exemplifies how nanotube-dependent gene transfer benefits cancers

Third, our bioinformatic analyses pipeline helped prioritize one borrowed gene, *CCDC88A*/GIV, from a list of several interesting candidates. We rationalized GIV because of its known pro-tumorigenic properties (outlined earlier in *Results*) and because we found it intriguing that the ER+ breast cancer cells typically silence this gene through alternative splicing (51), only to ‘borrow’ it back from MSCs later in the bone marrow niche (this work). We show transfer of GIV primarily as protein (not RNA). Because protein transfer is not driven by gradients and instead relies upon mechanisms of sorting and selective exchange, what governs selectivity of the borrowed proteins remains unclear. For GIV, we show that it binds and co-localizes with CX43 near gap junctions and within TNTs. The two proteins co-transfer from MSCs to cancer cells in ways that require functional gap junctions. Discovered originally as an actin binding/remodeling protein in breast cancers (50, 51) and found later to directly bind microtubules and localize to cell junctions under bioenergetic stress (57), both actin and microtubule filaments may serve as highways for transport of GIV between MSCs and cancer cells as described previously for motor proteins (15) within TNTs. Alternatively, because GIV associates with and regulates functions and/or localization of diverse cellular organelles (ERGIC-Golgi vesicles (93), exocytic vesicles (94), endosomes (94, 95), autophagosomes (96)) and cell surface receptors (e.g., EGFR, integrins, TLR4 etc.) (97), co-transport with such organelles or membrane proteins is also possible. While we did not resolve which of these mechanisms of transfer occur, we assessed the impact of GIV transfer.

By stably expressing GIV alone (among the list of borrowed candidates) in an ER+ breast cancer cell line, we recapitulated ∼20% of the gene expression patterns within the borrowed and intrinsic response components. Expression of GIV in ER+ breast cancer cells conferred resistance to clinical anti-estrogen drugs and promoted early survival and dissemination of circulating breast cancer cells. *CCDC88A*/GIV is also induced in ER+ breast cancer cells in contact cultures with fibroblasts, keeping with prior work demonstrating the role of GIV in cancer-associated fibroblasts in the dissemination of cancer (66). MSCs and other sources of carcinoma-associated fibroblasts in primary breast tumors potentially may donate GIV to ER+ breast cancer cells, priming cancer cells to survive in the circulation and disseminate to bone marrow. Given the short-lived duration of GIV in recipient breast cancer cells after separation from MSCs we observed in our studies (**Fig 4E**), ER+ breast cancer cells upon landing in the bone marrow may then need to re-establish contacts with MSCs in bone marrow to maintain expression of GIV. Considering recent revelations that GIV scaffolds a circuit for secretion coupled cellular autonomy (or growth signaling autonomy) in multicellular eukaryotic cells (32), it is possible that sharing of GIV between haves (stromal cells) and have nots (tumors that are originally GIV-negative) as ‘ready-to-use’ protein reflects a core biological phenomenon of sharing of resources during an urgent need for survival in growth factor restrictive environments. Transfer as protein bypasses the need for energy expenditure for protein translation and yet achieves higher levels of protein than possible through mRNA transfer; the latter is estimated to be never more than >1% of levels in donor cells(98). Although we found that GIV protein acquired during 3 days in contact culture was rapidly lost after returning cancer cells to monoculture, prolonged contact cultures (for months and years, as happens in patients) may allow transferred GIV to initiate the GIV↔STAT3 feedforward loop shown to enhance its own transcription (99) and, thereby, sustain its levels in cells. If so, GIV may promote further ‘social’ interactions and ‘exchanges’ with bone marrow MSCs through augmentation of TNT formation to facilitate its ‘spread’ across tumor masses as tumors grow. This is not entirely unexpected given the distinct microtubule and actin-binding modules present in GIV and its known functions in vesicle transport, all aspects that are critical for active transport via TNTs (30, 54). We also noted that GIV mRNA was elevated in contact cultures, suggesting that acquisition of some GIV as mRNA or its induction as an intrinsic tumor cell response to the ‘borrowed’ proteins likely occurs. We conclude that intercellular protein transport is a major way for cells to exchange GIV as an essential commodity within a heterotypic population of haves and have nots.

#### Study limitations

A limitation of this study is the use of a conditioned medium from MSCs rather than contact-free co-cultures with separation of cell types in transwells. Although such a setup may have enabled assessment of bidirectional crosstalk in which ER+ tumor cells could initiate or perpetuate the effects of MSCs on cancer cells, transwell setups present substantial technical challenges and risk a major confounding factor. For example, it is challenging to collect sufficient cells for RNA sequencing and proteomics from transwells, and prolonged (72 h) cultures risk the formation of gradients of secreted growth factors and/or chemokines that could trigger migration across the insert and direct contact between cell types. Also, we did not pursue detailed characterization of all that is transported. Some genes are likely to be transported from cancer cells to MSCs; we did not evaluate this here. Contact culture induced tolerant immunogenic signals in cancer cells; how these signals impact immune evasion by cancer cells and/or resistance to immune checkpoint blockade therapies will require further studies in immunocompetent mice. TNT-mediated tumor-CAF exchanges also shape tumor cell behaviors (65, 100); dedicated studies are required to further investigate the shared and distinct patterns between these (and potentially others, e.g., tumor-myeloid cell) forms of cell-cell communication. Further studies also are needed to establish mechanisms initiating and sustaining TNT communication between cancer and stromal cells and if the nature and extent of the TNT-facilitated exchange described here also occur in more complex systems, such as patient-derived organoids cultured in 3D under near-physiologic conditions in the presence of immune and non-immune cells. Furthermore, we did not distinguish which RNAs and proteins presumed to be ‘borrowed’ from MSCs by ER+ breast cancer cells are directly transferred as opposed to reflecting secondary changes induced by a subset of the borrowed molecules. Regardless of the relative contributions of these subsets, the induced transcriptome-proteome carries meaningful information regarding tumor growth, resistance, and relapse.

In conclusion, this work provides insights as to how close contact interactions and intercellular communication networks through RNA/protein transfer between ER+ breast cancer cells and MSCs reprogram tumor cells to more aggressive states. Based on the omics-based revelations and experimental evidence (this work) and phenotypic characterization of MSC-tumor contact culture we published earlier (68), we conclude that TNT-based cell-cell communication in the bone marrow niche (or tumor-stromal or tumor-immune cell communication in any other tissue) may provide the same to cancer cells that nanotube-based communication provides to the bacteria within biofilms; a nutrition- and metabolic stress-free haven to promote sustained survival in new environments. Blocking or redirecting these communication networks, or components transferred through these networks, offers promising opportunities to prevent or cure breast cancer metastasis to bone.

## Materials and Methods

### Sex as a Biologic Variable

All animal experiments used only female mice because 99% of breast cancers occur in women.

### RNA sequencing and differentially expressed gene (DEG) analysis

Genes with log fold change >=2 and a p adjusted value < 0.05 were identified and rank ordered as differentially expressed genes (DEGs) (**Supplemental Information 5**). Besides the datasets generated in this work, we leveraged several publicly available datasets. A complete inventory of these datasets and their nature, composition, and source is presented in **Supplemental Information 7**.

### Tandem Mass Tag™ (TMT) proteomics and analyses

We deposited mass spectrometry proteomics data in the ProteomeXchange Consortium via the PRIDE(101) partner repository with the dataset identifier (**PXD039860**). A list of differentially expressed proteins is provided in **Supplemental Information 6**.

### Study Approval

The University of Michigan IACUC approved all animal procedures under protocol number PRO00010534.

### Data Availability

Data are available via ProteomXchange with identifier PDX039860 and Gene expression Omnibus with the identifier GSE224322. The computational analyses can be found at https://github.com/sinha7290/cx43. Values for all data points in graphs are reported in the Supporting Data Values file.

**Supplemental Material and Methods** contains further information about experimental and computational procedures.

## ACKNOWLEDGEMENTS

The authors acknowledge funding from United States National Institute of Health grants R01CA238042 (P.G. and G.D.L.); R01CA100768, R01CA160911 (P.G.); R50CA221807 (K.E.L) and U01CA210152, R01CA238023, R33CA225549, and R37CA222563 (G.D.L). We also acknowledge support from the Padres Pedal the Cause #PTC2021 and the Torey Coast Foundation, La Jolla (P.G) and the W.M. Keck Foundation (G.D.L). SS was supported in part by the American Association of Immunologists’ (AAI) Intersect Fellowship Program for Computational Scientists and Immunologists. We thank Guillaume Castillon (UC San Diego Electron Microscopy Core Facility) for technical and logistical support and Debashis Sahoo (UC San Diego) for access to the computational platform, Hegemon. This manuscript includes data generated at the UC San Diego Institute of Genomic Medicine (IGM) using an Illumina NovaSeq 6000 that was purchased with funding from a National Institutes of Health SIG grant (#S10 OD026929). The manuscript also includes data generated through the University of Michigan Advanced Genomics, Flow Cytometry, and Pre-Clinical Imaging shared resources supported in part by the University of Michigan Comprehensive Cancer Center support grant P30CA046592.

## AUTHOR CONTRIBUTIONS

S.S, supervised by P.G, carried out and analyzed all electron microscopy, RNA seq, TMT proteomics and computational analyses showcased in this work. Su.R. carried out protein purification and GST pulldown assays. B.W.C, A.P.F., S.C., Sh.R., and J.M.B performed mouse experiments and cell-based assays. K.E.L. provided reagents. B.W.C. S.S, P.G. and G.D.L. assembled the figures. K.E.L., Su.R., S.S., P.G., and G.D.L. helped write methods and edit the manuscript. G.D.L. and P.G conceptualized, designed, supervised, and analyzed the experiments. K.E.L, S.S. P.G and G.D.L wrote the manuscript. All authors approved the manuscript prior to submission.

