## Supplementary Online Materials for "Breast Cancers That Disseminate to Bone Marrow Acquire Aggressive Phenotypes through CX43-related Tumor-Stroma Tunnels"


**Authors and affiliations**

Saptarshi Sinha[1], Brennan W. Callow[2], Alex P. Farfel[2], Suchismita Roy[1], Siyi Chen[2], Shrila Rajendran[2], Johanna M. Buschhaus[2,3], Celia R. Espinoza[1], Kathryn E. Luker[2,4], Pradipta Ghosh[1,5,6,7*], Gary D. Luker[2,3,4*]

[1]Department of Cellular and Molecular Medicine, School of Medicine, University of California San Diego, La Jolla, CA, USA

[2] Center for Molecular Imaging, Department of Radiology, University of Michigan, Ann Arbor, MI, USA

[3]Department of Biomedical Engineering, University of Michigan, Ann Arbor, MI, USA

[4]Biointerfaces Institute, University of Michigan, Ann Arbor, MI, USA

[5]Moores Comprehensive Cancer Center, [6]Department of Medicine, [7]School of Medicine, and Veterans Affairs Medical Center, University of University of California San Diego, La Jolla, CA

*Corresponding authors:

Pradipta Ghosh, Departments of Medicine and Cellular and Molecular Medicine, University of California San Diego, 9500 Gilman Drive (MC 0651), George E. Palade Bldg, Rm 232, 239, La Jolla, CA 92093; Phone: 858-822-7633.

Gary D. Luker, Departments of Radiology and Biomedical Engineering, A524 BSRB, 109 Zina Pitcher Place, Ann Arbor, MI 48109-2200. Phone: 734-764-2890.

### INVENTORY OF SUPPLEMENTARY ONLINE MATERIALS

- **Supplementary Figures and Legends (7)**
- **Supplementary Methods**
- **Supplementary References**
- **Additional Supplementary Materials** (Uploaded separately as Excel sheets)
  1. **Supplemental information 1:** Full catalog of all upregulated genes and proteins in contact culture
  2. **Supplemental information 2:** Full catalog of all downregulated genes and proteins in contact culture
  3. **Supplemental information 3:** Edge and node list of the entire PPI network and various subnetworks
  4. **Supplemental information 4:** Full catalog of all genes presented in Figure 6A.
  5. **Supplemental information 5:** Inventory of differentially expressed genes presented in Figure 1C.
  6. **Supplemental information 6:** Inventory of differentially expressed proteins presented in Figure 1C.
  7. **Supplemental information 7:** Inventory of publicly available datasets used in this study.

### SUPPLEMTARY FIGURES AND LEGENDS

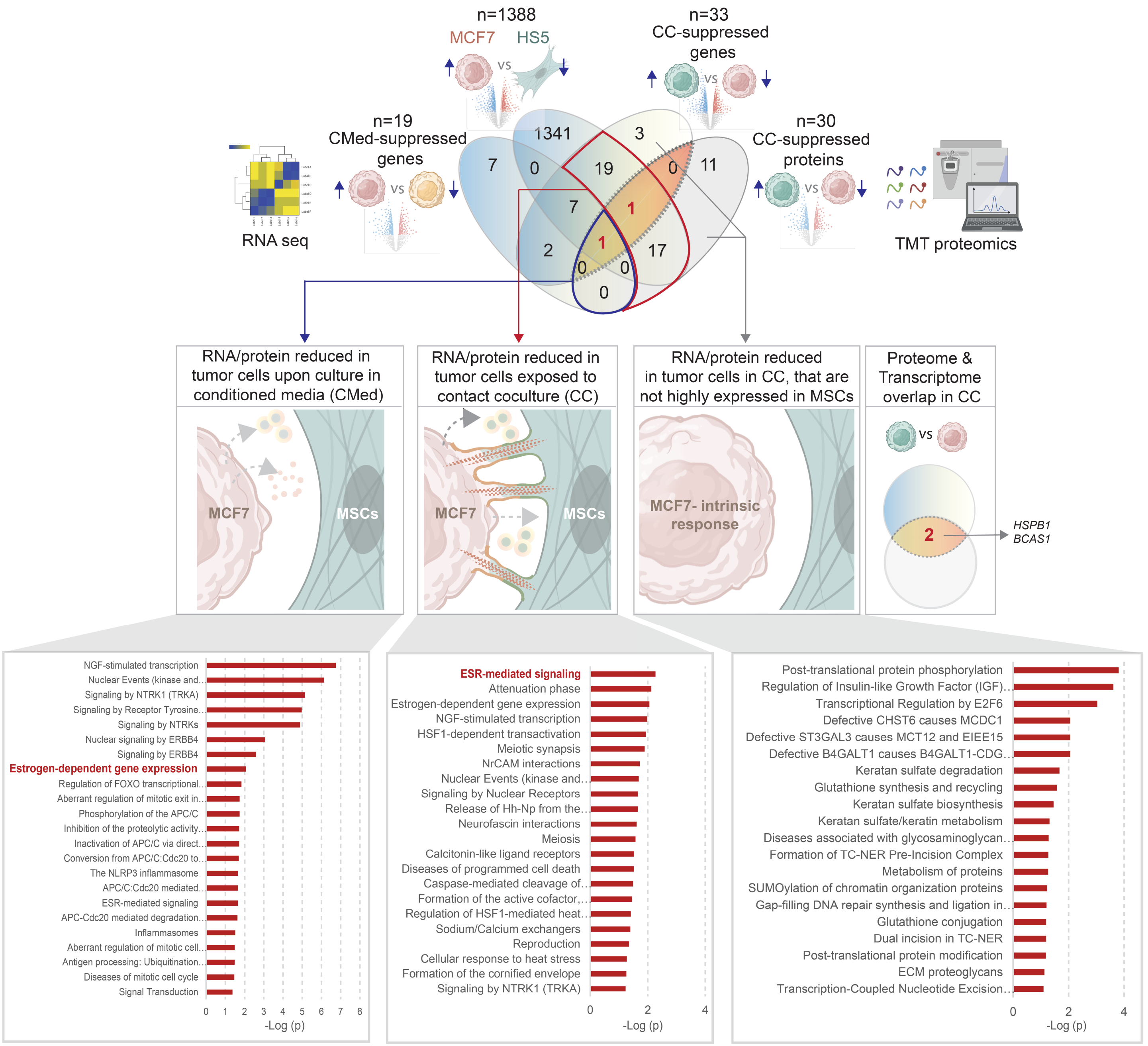


**Supplementary Figure 1**. **Borrowed (from MSCs) and tumor-intrinsic gene induction patterns carry distinct meaning**. Pathway enrichment analyses for genes/proteins induced during contact co-culture (CC; A) for a subset of 39 genes induced in CC which were also detected via proteomics (CC; B), for genes/proteins induced upon exposure to conditioned media (CMed; C) and for genes/proteins induced in MCF7 tumor cells that are not highly expressed in HS5 MSCs, and, hence, presumed to be tumor cell intrinsic responses (D).

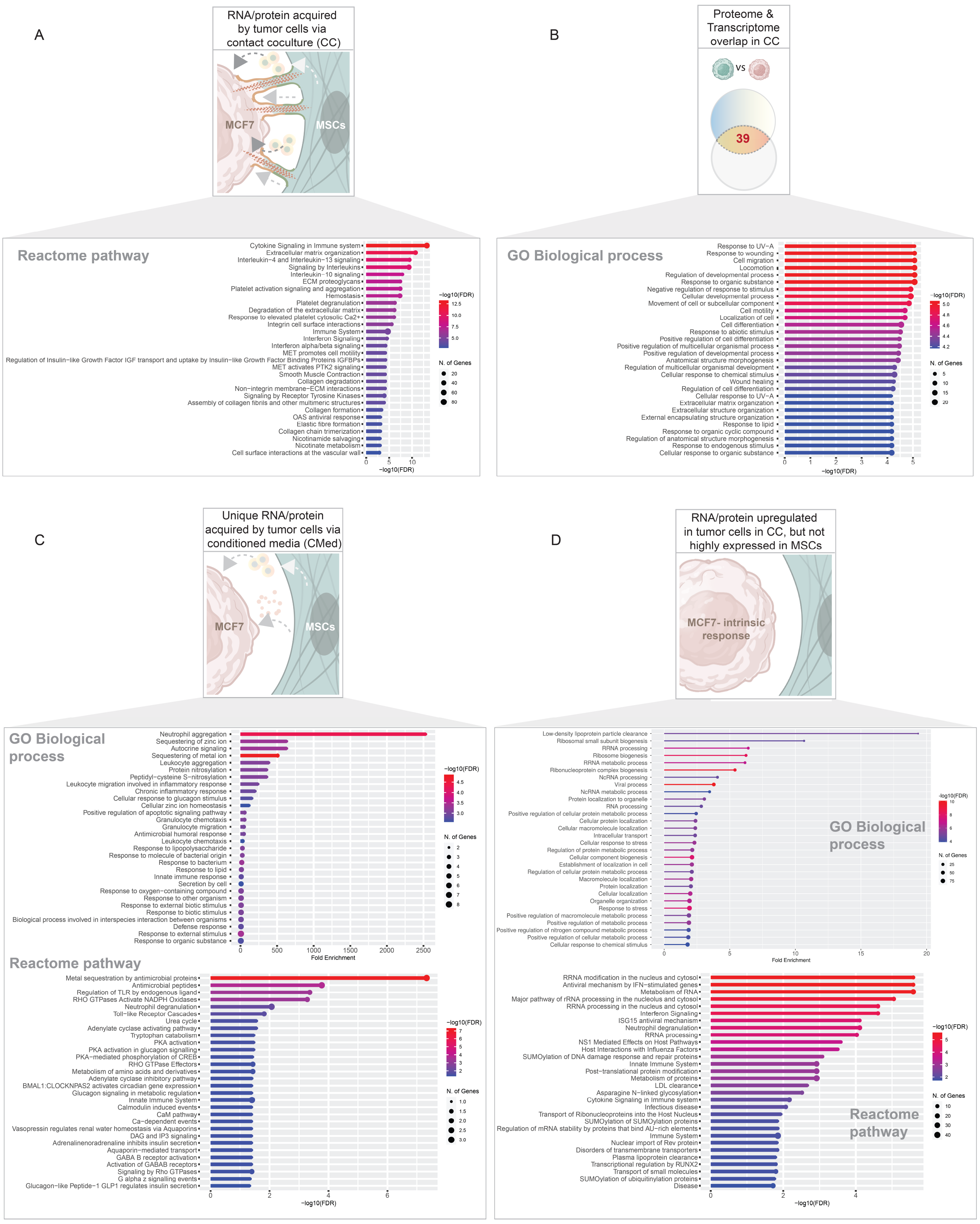


**Supplementary Figure 2: Estrogen-dependent signaling is suppressed in ER-positive breast cancer cells after co-culture with bone marrow mesenchymal stem cells.**

*Top*: Venn diagram depicts the outcome of multiple DEG analyses between different comparator groups that catalogs the number of suppressed genes or proteins in contact co-culture (CC) or in conditioned media (CMed) and the overall suppressed genes in MCF7 cells compared to the HS5 bone marrow MSCs. Red = 2 uniquely downregulated transcripts in tumor cells in contact co-culture that were also identified by proteomics. Genes/proteins that are suppressed in MCF7 tumor cells in coculture with HS5 MSCs are binned into 3 categories (connected by arrows) based on the likely mechanisms for such induction.

*Bottom*: Reactome pathways enriched in the downregulated genes/proteins identified in each category.

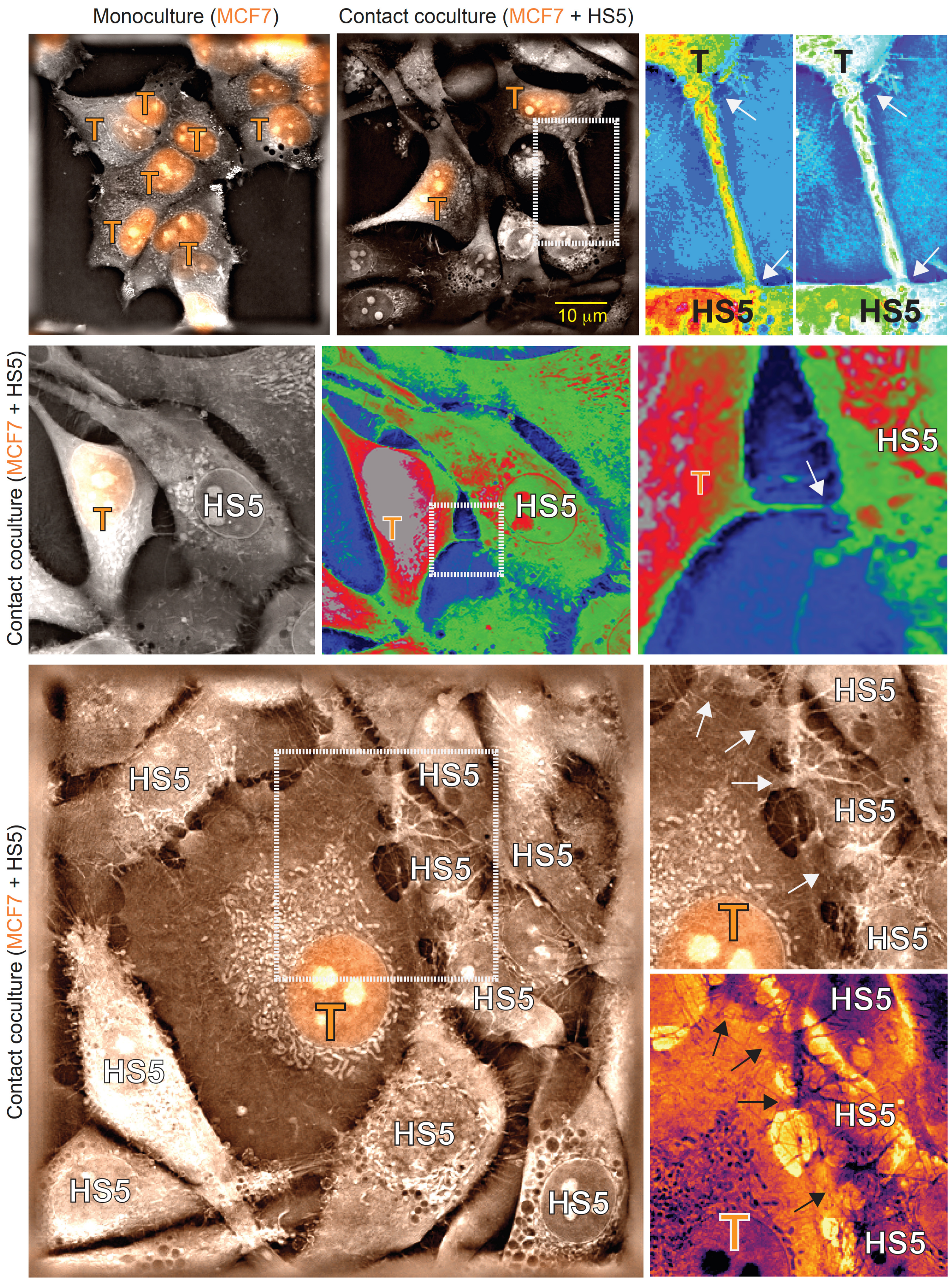


**Supplementary Figure 3: MCF7 cells form tunneling nanotubes with HS5 cells in co-culture.** Live-cell interferometry microscopy images of MCF7 cells stably transduced with nuclear mCherry (denoted as T) in monoculture (upper left) or co-cultures with HS5 cells (all other panels). We pseudocolored selected boxed aspects of images to emphasize tunneling nanotubes (white arrows) between MCF7 and HS5 cells. Diameter of tunneling nanotubes (TNTs) is larger at the MCF7 side of the connection with HS5 cells, suggesting tube formation originated from MCF7 cells. Scale bar = 10 µm.

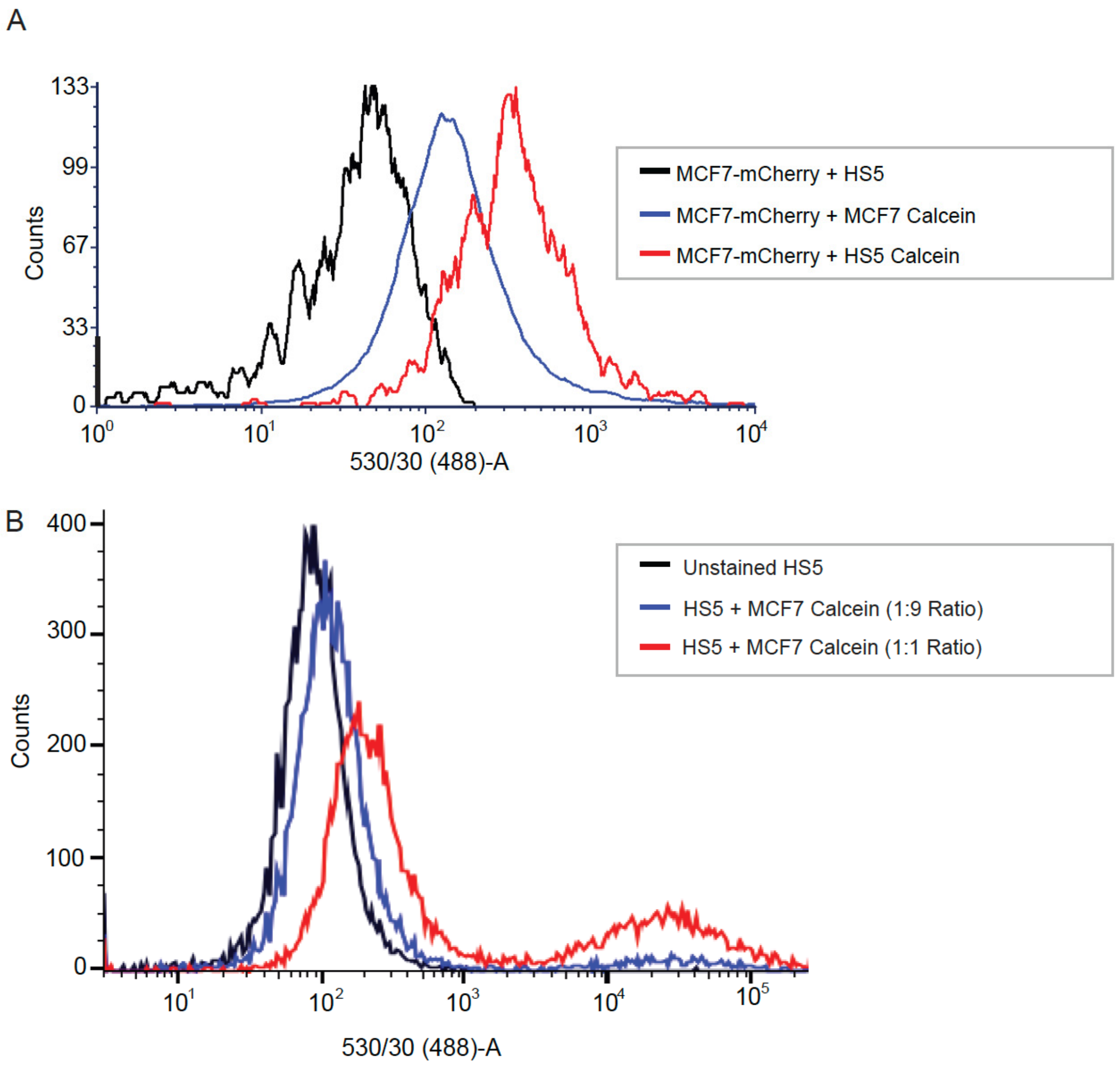


Supplementary Figure 4: **MCF7 and HS5 cells establish functional structures for bidirectional intercellular communication. A.** MCF7 or HS5 cells labeled with fluorescent dye calcein AM (donors) were co-cultured with MCF7 cells that stably expressed mCherry (recipient) for 3 d prior to the analysis of transferred calcein by flow cytometry. Both HS5 and MCF7 cells transferred calcein to recipient MCF7 cells (the latter were marked by mCherry). However, HS5 cells transferred markedly more calcein to MCF-mCherry cells than MCF7 donor cells. Transfer of calcein from HS5 to MCF7 recipient cells demonstrates that these cells form functional intercellular contacts that allow exchange of cytosolic small molecules. **B.** We also labeled parental MCF7 cells with calcein (donor) before co-culturing with HS5 MSCs (recipients) at ratios of 1:9 or 1:1 HS5 to MCF7 cells. The 1:9 ratio of HS5 to donor cells is the opposite of the standard conditions used for our other co-cultures (9:1 MSCs to breast cancer cells). HS5 cells stably expressed mCherry. Flow plot gated on HS5-mCherry cells shows transfer of calcein from MCF7 to HS5 cells with greater amounts transferred in the 1:1 co-culture condition.

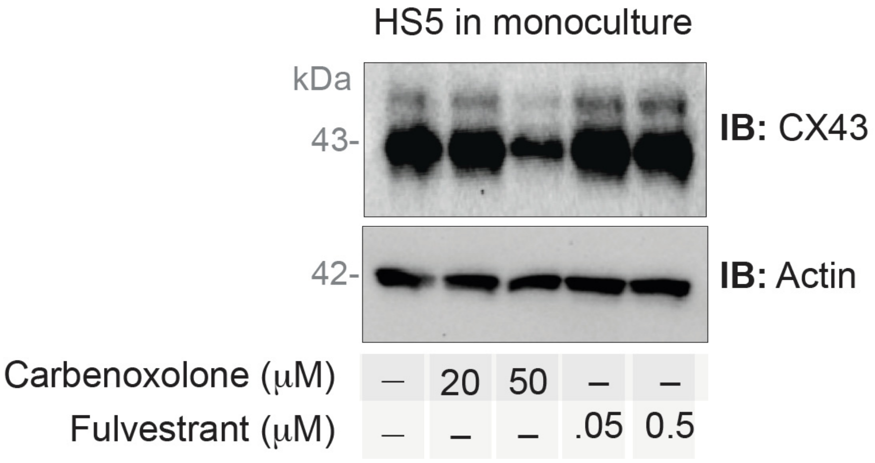


**Supplementary Figure 5: Effect of carbenoxolone or fulvestrant on Cx43 expression in HS5 cells.** Monocultures of HS5 cells were treated with the indicated concentrations of Carbenoxolone or Fulvestrant prior to lysis. Equal aliquots of lysates were analyzed for Cx43 or actin (as a loading control) by immunoblotting (IB).

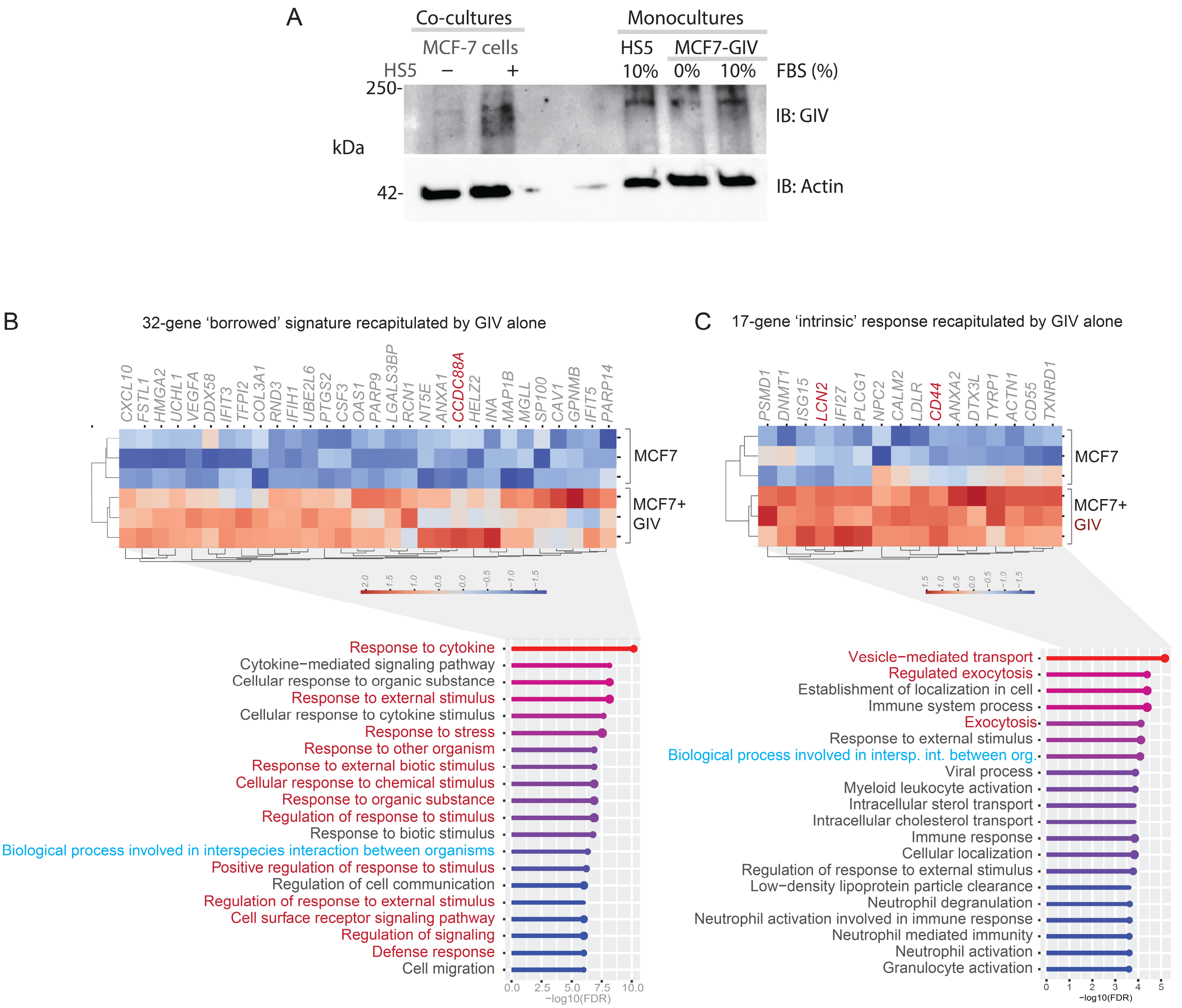


**Supplementary Figure 6. Stable expression of GIV in MCF7 cells (A) recapitulates components of the borrowed (B) and tumor-intrinsic (C) gene signature from co-cultures of breast cancer cells with MSCs.** Immunoblots (A) show that the levels of exogenous expression of GIV in MCF7-GIV cells are comparable to the levels of expression of GIV in the same cells after 72 h of contact culture with HS5 cells. Heatmaps display the hierarchical unsupervised clustering of MCF7 and MCF7-GIV cells by z-score normalized gene expression patterns for the subset of genes that are induced among the borrowed (B) and intrinsic (C) signatures. Reactome pathway analysis of these genes are displayed underneath each heatmap. Red font highlights genes of interest indicative of cancer cell stemness (top) or biological processes (bottom) associated with cellular response to stimuli or stress (B) or membrane transport and trafficking (C). Blue fonts highlight processes related to cell-cell interactions.

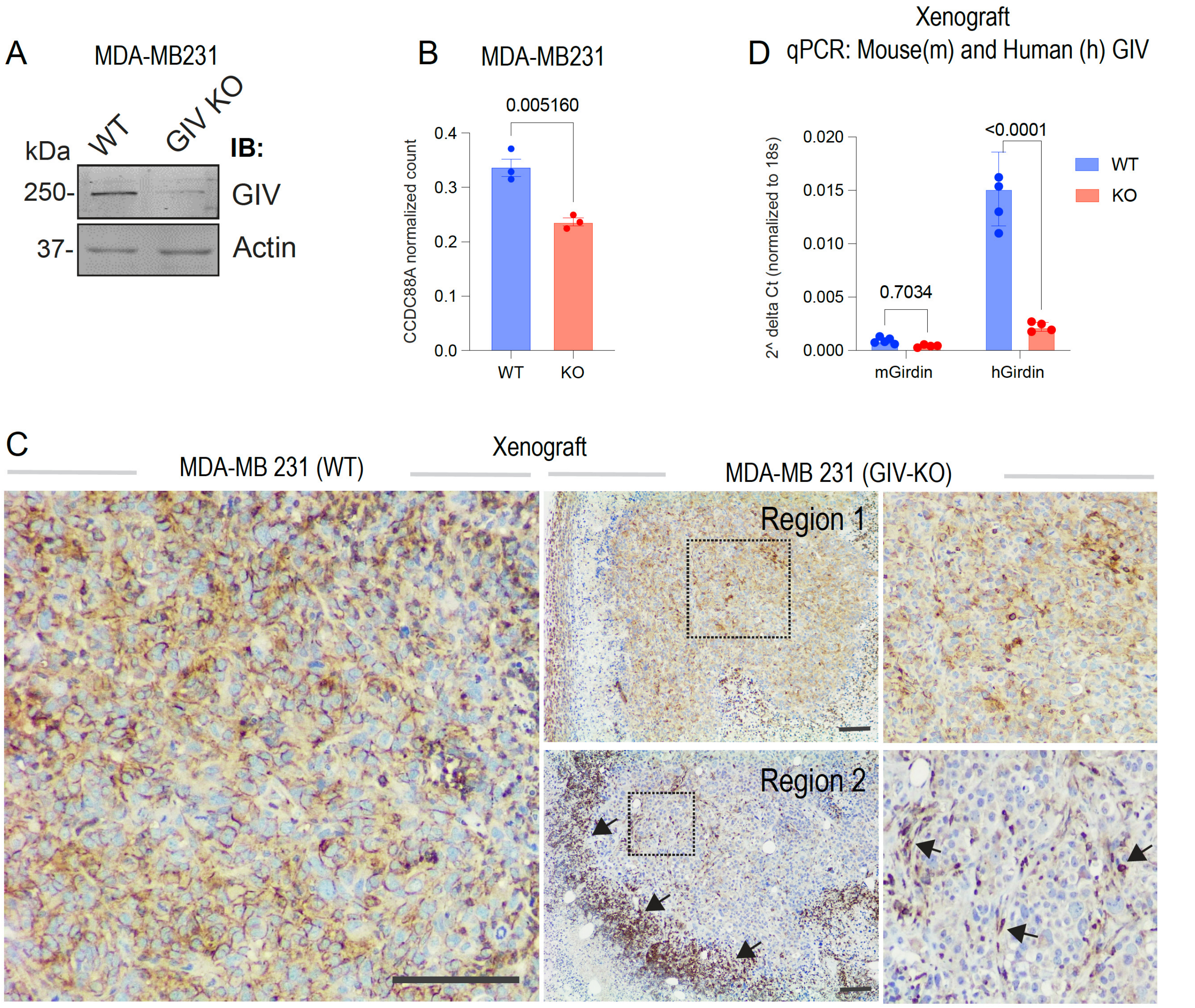


**Supplementary Figure 7. Cancer-associated fibroblasts could serve as 'donors' of GIV in primary tumors.** A-B) CRISPR-mediated GIV depletion in MDA-MB-231 cells: immunoblotting (A) and estimation of transcript counts by RNA Seq(1) [GSE215822] (B). C) Tumors implanted in the mammary fat pads of NSG mice were analyzed for GIV by IHC (Millipore; Abt80). Scale bar = 200 μm. GIV+ tumor regions were observed in GIV-KO tumors, surrounded by GIV+ stromal cells (arrows). D) qPCR confirming the absence of mouse GIV and continued depletion of human transcripts.

### Supplemental Materials and Methods

#### Cell culture

We purchased MCF7 (HTB-22) and T47D (HTB-133) human ER+ breast cancer cell lines, Cos7 cells, MDA-MB-231 human breast cancer cells, and HS5 (CRL-11882) and HS27a (CRL-2496) human bone marrow MSC lines from the American Type Culture Collection (American Type Culture Collection, ATCC). We thank Dr. James Rae (University of Michigan) for providing human HCC1428 ER+ breast cancer cells and Dr. Daniel Hayes (University of Michigan) for immortalized human mammary fibroblasts. We previously have described MDA-MB-231 cells with knockout (KO) of GIV by CRISPR/Cas9 and stable expression of click beetle green luciferase in parental MDA-MB-231 and MDA-MB-231 GIV KO cells (2). We cultured HS5, MCF7, T47D, MDA-MB-231, human mammary fibroblasts, and Cos7 cells in DMEM (#11995, Gibco, Thermo Fisher) with 10% fetal bovine serum (FBS) (HyClone, ThermoFisher Scientific), 1% GlutaMAX (#35050, Gibco), and 1% Penicillin-Streptomycin (P/S, #15140, Gibco). We cultured HCC1428 and HS27a cells in RPMI (#11875, Gibco) with 10% FBS, 1% P/S, 1% Sodium Pyruvate (#11360070, Gibco), 1% HEPES (#15630080, Gibco), and 2500 mg/L glucose (#A24940, Gibco). We added 50mg/L Plasmocin™ prophylactic (Invivogen, San Diego, CA, USA) to both DMEM and RPMI media. We maintained cells as previously described (3). We performed mycoplasma testing and authenticated cell lines with short tandem repeats analysis upon initial passaging. We removed cell lines from culture after three months and replaced them with new cells from previously frozen stocks.

#### Vectors and stable cell lines

We previously described MCF7 cells stably transduced with click beetle green luciferase (MCF7-CBG) (4). For stable RNA interference against GIV, we transduced MCF7 and HS5 cells with shRNA against *CCDC88A* (clone TRCN0000129915, Millipore Sigma), preparing lentiviruses as described previously (5). We used puromycin to select stably transduced cells and performed experiments with batch populations. To stably express full-length GIV in MCF7-CBG cells, we used a PiggyBac transposon with a CAG promoter driving co-expression of GIV and hygromycin linked by a P2A site (VectorBuilder). We transfected cells with a 3:1 ratio of GIV transposon to Super PiggyBac Transposase (Systems Biosciences) using Fugene-HD transfection reagent (Promega) according to the manufacturer's protocol. Two days after transfection, we selected cells with 150 μg/ml hygromycin B (ThermoFisher Scientific). After two weeks of culture with hygromycin B, we returned surviving cells to normal growth medium. We transduced MCF7-CBG cells with a lentiviral vector expressing mCherry with a nuclear localization sequence as we reported previously (6).

#### Co-cultures of breast cancer cells and MSCs

For co-culture assays, we seeded 12,000 total cells per $cm^2$ in 35 mm dishes with a ratio of 1:9 ER+ breast cancer cells to MSCs. One day after seeding, we washed cells once with phosphate buffered saline (PBS) and then changed medium to phenol-red free DMEM (#A14430-01, Gibco), supplemented with 1% FBS, 1% P/S, 1% Glutamax, 1% sodium pyruvate, 10 nM β-estradiol (#E2758, Sigma-Aldrich, Millipore Sigma), and 5 mM glucose (7). We cultured cells in low serum (1%), low glucose (4 mM), and 10 nM estrogen (LG/LF) medium for

3 days before analyzing experimental endpoints. We performed assays with conditioned medium as described previously (8).

**Interferometry Microscopy**

We cultured MCF7 cells alone (monoculture) or in co-culture with HS5 or HS27a cells on 35 mm dishes with glass bottoms (Cellvis) as described above in the section for co-cultures of breast cancer cells and MSCs. For these studies, we used MCF7 cells stably expressing nuclear localized mCherry to distinguish these cells from MSCs. We imaged cells using a Nanolive 3D Cell Explorer microscope (Nanolive) equipped with a 60X objective. The instrument has a stage top incubator for live cell imaging. We collected images from >15 randomly selected fields per sample (3 biologic replicates per condition), using fluorescence from mCherry to identify MCF7 cells. A person blinded to experimental conditions quantified numbers of tunneling nanotubes (TNTs) connecting MCF7 to MCF7 cells or MCF7 cells to MSCs in collected images.

**RNA sequencing and differentially expressed gene (DEG) analysis**

We separated cancer cells from stromal cells with human EpCAM (CD326) immunomagnetic beads (#130-061-101, Miltenyi) for RNA sequencing as described previously (9). RNA extraction was done using a kit (R2052, Zymo Research) as per manufacturer's protocol. RNA sequencing libraries were generated using the Illumina TruSeq Stranded Total RNA Library Prep Gold with TruSeq Unique Dual Indexes (Illumina). Samples were processed following the manufacturer's instructions except modifying RNA shear time to five minutes. Resulting libraries were multiplexed and sequenced with 100 basepair (bp) Paired End (PE100) to a depth of approximately 25-40 million reads per sample on an Illumina NovaSeq 6000. Samples were demultiplexed using bcl2fastq v2.20 Conversion Software (Illumina). RNASeq data were processed using kallisto (version 0.45.0). Gene-level TPM values and gene annotations were computed using a custom Perl Script. We deposited RNA sequencing data to GEO under accession number GSE224322. Differentially expressed genes from the pairs of experimental conditions are identified using DESeq2 program in R. Genes with log fold change >=2 and a p adjusted value < 0.05 were identified and rank ordered as differentially expressed genes (DEGs) (**Supplemental Information 5**). Besides the datasets generated in this work, we leveraged several publicly available datasets. A complete inventory of these datasets and their nature, composition, and source is presented in **Supplemental Information 7**.

**Hierarchical clustering of RNA seq readouts**

Expression patterns of the 'borrowed' core set of 39 genes that are highly expressed in MCF7 cells after coculture with HS5 cells (**Fig 1**) were used to arrange all the samples in the datasets in an unsupervised way based on their z normalized expression using the seaborn clustermap package (v 0.12) in python.

**Tandem Mass Tag™ (TMT) proteomics and analyses**

MCF7 cells from monoculture and post coculture with HS5 cells were subsequently processed for TMT proteomics using LUMOS Orbitrap-Fusion analyzer. Samples were processed at the UC San Diego Biomolecular and Proteomics Mass Spectrometry Core Facility (https://bpmsf.ucsd.edu/). For protein digestion, 3 mls of 6 Molar Guanidine solution was added to cell pellet and mixed. The samples were then boiled for 5 minutes followed by 5 minutes cooling at room temperature. The boiling and cooling cycle was repeated for a total of 3 cycles. The proteins were precipitated with addition of methanol to a final volume of 90% followed by vortex and centrifugation at maximum speed on a benchtop microfuge (4000 rpm) for 20 minutes. The soluble fraction was removed by flipping the tube onto an absorbent surface and tapping to remove any liquid. The pellet was suspended in 4ml of 8 M Urea made in 100mM Ammonium Bicarbonate. TCEP was added to final concentration of 10 mM and chloro-acetamide solution was added to final concentration of 40 mM with vortexing for 5 minutes. 3 volumes of 50mM ammonium bicarbonate were added to the sample to reduce the final urea concentration to 2 M. Trypsin was added at a 1:50 ratio of trypsin to sample and incubated at 37C for 48 hours. The solution was then acidified using trifluoroacetic acid (TFA) (0.5% TFA final concentration) and mixed. Samples were desalted using 100 mg C18-SPR (Waters) as described by the manufacturer's protocol. The peptide concentration of each sample was measured using a BCA assay. 100 µg of each sample were then labeled using TMT10 (as suggested in the manufacturer protocol, ThermoFisher Scientific) for one hour and quenched using hydroxylamine. The samples were then pooled and dried. High pH fractionation: Pierce™ High pH Reversed-Phase Peptide Fractionation Kit (Pierce™ High pH Reversed-Phase Peptide Fractionation Kit Catalog number: 84868) was used. The fractionation protocol was performed as described by the manufacturer's kit with the exception that 12 fractions were generated. LC-MS-MS: Each fraction was analyzed by ultra-high-pressure liquid chromatography (UPLC) coupled with tandem mass spectroscopy (LC-MS/MS) using nano-spray ionization. The nano-spray ionization experiments were performed using an Orbitrap fusion Lumos hybrid mass spectrometer (Thermo) interfaced with nano-scale reversed-phase UPLC (Thermo Dionex UltiMate™ 3000 RSLC nano System) using a 25 cm, 75-micron ID glass capillary packed with 1.7-µm C18 (130) BEHTM beads (Waters corporation). Peptides were eluted from the C18 column into the mass spectrometer using a linear gradient (5–80%) of acetonitrile (ACN) at a flow rate of 375 µl/min for 120 min. The buffers used to create the ACN gradient were the following: Buffer A (98% H2O, 2% ACN, 0.1% formic acid) and Buffer B (100% ACN, 0.1% formic acid). Mass spectrometer parameters are as follows: an MS1 survey scan using the orbitrap detector (mass range (m/z): 400-1500 (using quadrupole isolation), 60000 resolution setting, spray voltage of 2200 V, ion transfer tube temperature of 275 C, AGC target of 400000, and maximum injection time of 50 ms) was followed by data dependent scans (top speed for most intense ions, with charge state set to only include +2-5 ions, and 5 second exclusion time, while selecting ions with minimal intensities of 50000 at in which the collision event was carried out in the high energy collision cell (HCD Collision Energy of 38%) and the first quadrupole isolation window was set at 0.7 (m/z). The fragment masses were analyzed in the Orbi-trap mass analyzer with mass resolution setting of 15000 (With ion trap scan rate of turbo, first mass m/z was 100, AGC Target 20000 and maximum injection time of 22ms). Peptides are identified and mapped using Peaks Studio X (Bioinformatics Solutions Inc.). Intensity ratio of each identified protein in MCF7 cell monoculture and post coculture with HS5 cells has been identified and selected if the significance score >20. The mass spectrometry proteomics data have been deposited to the ProteomeXchange Consortium via the PRIDE (10) partner repository with the dataset identifier (**PXD039860**). A list of differentially expressed proteins is provided in **Supplemental Information 6**.

**Protein-protein interaction network construction**

The Human Protein-Protein Interaction (PPI) data was retrieved from the STRING database (https://string-db.org/). To mitigate false positive interactions and enhance confidence, interactions with a combined score (generated by the STRING database) above 650 were used to construct the network; this cutoff was determined based on the highest possible combined score, ensuring coverage of at least one interaction associated with each protein in our seed list. Subsequently, a specific PPI network was constructed as a subgraph encompassing all necessary nodes to connect the seed proteins via the shortest paths within the network. Utilizing this algorithm, we devised a tailored PPI network linked to the seed protein list.

**StepMiner analysis**

StepMiner is an algorithm that identifies step-wise transitions using step function in time-series data (11). StepMiner undergoes an adaptive regression scheme to verify the best possible up and down steps based on sum-of-square errors. The steps are placed between time points at the sharpest change between expression levels, which gives us the information about timing of the gene expression-switching event. To fit a step function, the algorithm evaluates all possible steps for each position and computes the average of the values on both sides of a step for the constant segments. An adaptive regression scheme is used that chooses the step positions that minimize the square error with the fitted data. Finally, a regression test statistic is computed as follows:

$$F\ stat = \frac{\sum_{i=1}^{n}(\widehat{X_i} - \bar{X})^2/(m-1)}{\sum_{i=1}^{n}(X_i - \widehat{X_i})^2/(n-m)}$$

Where $X_i$ for $i = 1$ to $n$ are the values, $\widehat{X_i}$ for $i = 1$ to $n$ are fitted values. m is the degrees of freedom used for the adaptive regression analysis. $\bar{X}$ is the average of all the values: $\bar{X} = \frac{1}{n} * \sum_{j=1}^{n} X_j$. For a step position at k, the fitted values $\widehat{X_i}$ are computed by using $\frac{1}{k} * \sum_{j=1}^{n} X_j$ for $i = 1$ to $k$ and $\frac{1}{(n-k)} * \sum_{j=k+1}^{n} X_j$ for $i = k + 1$ to $n$.

**Composite gene signature analysis using Boolean Network Explorer (BoNE)**

Boolean network explorer (BoNE) (12) provides an integrated platform for the construction, visualization and querying of a gene expression signature underlying a disease or a biological process in three steps: First, the expression levels of all genes in these datasets were converted to binary values (high or low) using the StepMiner algorithm. Second, Gene expression values were normalized according to a modified Z-score approach centered around *StepMiner* threshold (formula = (expr - SThr)/3*stddev). Third, the normalized expression values for every gene were added together to create the final composite score for the gene signature. These composite scores represent the overall activity or state of biological pathways associated with the genes and, hence, can identify differences between control and query groups *within* any given dataset. However, composite scores can neither be directly compared between various gene signatures on a dataset (because they are not normalized according to the number of genes in each signature), nor can the same signature be compared across datasets (which are individually normalized according to intra-dataset sample distribution or have other inherent differences). The samples were ordered based on the final signature score. Classification of sample categories using this ordering is measured by ROC-AUC (Receiver Operating Characteristics Area Under The Curve) values. Welch's Two Sample t-test (unpaired, unequal variance (equal_var=False), and unequal sample size) parameters were used to compare the differential signature score in different sample categories. Violin, swarm and bubble plots are created using python seaborn package version 0.10.1. Pathway enrichment analyses for genes were carried out via the reactome database (https://reactome.org/) and algorithm (13). Violin plots are created using python seaborn package version 0.10.1.

**Measurement of classification strength or prediction accuracy**:

To measure the strength of classification or prediction accuracy, Receiver Operating Characteristic (ROC) curves were generated. These curves illustrate the diagnostic ability of a binary classifier system (e.g., high vs. low gene expression levels) as its discrimination threshold is adjusted along with the sample order. ROC curves plot the True Positive Rate (TPR) against the False Positive Rate (FPR) at various threshold settings. The Area Under the Curve (AUC) quantifies the probability that a classifier will correctly rank randomly chosen samples into two groups. Alongside ROC AUC, other classification metrics such as accuracy ((TP + TN)/N; TP: True Positive; TN: True Negative; N: Total Number), precision (TP/(TP+FP); FP: False Positive), recall (TP/(TP+FN); FN: False Negative), and f1 score (2 * (precision * recall)/(precision + recall)) were computed. The Python Scikit-learn package was used to calculate the ROC-AUC values.

**Survival outcome analyses**

Kaplan-Meier (KM) analyses were done for gene signature identified from transcriptome and proteome overlap. The high and low groups were separated based on *StepMiner* threshold on the composite score of the gene expression values in the associated cohort. The threshold for separating high and low groups varies between cohorts and depends on the gene signature composite score distribution within the patients. The statistical significance of KM plots was assessed by log rank test. Kaplan-Meier analyses were performed using lifelines python package version 0.14.6. The dataset we used (GSE25066) is comprised of Her2-negative, ER-status annotated samples, the majority of which belong to the multicenter I-SPY 1 TRIAL. The latter evaluated patients with ≥ 3 cm tumors using early imaging and molecular signatures with outcomes of pathologic complete response (pCR) and relapse free survival (RFS) (14). This multicenter cohort is also notable for being comprised of 90% intermediate or high-grade cancers and 91% high risk by the 70-gene profile, which helps enrich for poor outcome. The I-SPY 1 population is enriched for tumors with a poor prognosis but is still heterogeneous in terms of rates of pCR and RFS. The cohort has a total of 508 patients; however, for stratifying patients based on their ER-status, we excluded 6 patients who did not have such information (n = 502; 2B-C). We also carried out subset analyses in which we selected treatment insensitive patients who had residual disease (RD) (n = 189; 2D) or treatment sensitive disease that showed pathologic complete response (pCR; n = 46). The second cohort we used is the ER-status annotated, Her2-negative population within the well-known METABRIC dataset (15) (https://ega-archive.org/studies/EGAS00000000083).

**Flow cytometry for intercellular transfer of calcein AM**

To measure transfer of dye between cells by intercellular connections, we incubated MCF7 or HS5 cells with 10 μM calcein AM (C3099, Thermo Fisher Scientific) in LG/LF medium for one hour. We then washed cells three times with PBS and seeded labeled MCF7 or HS5 cells with MCF7-mCherry cells at a 9:1 ratio in 10 cm dishes using the cell density described above for co-culture assays. After three days in co-culture, we analyzed amounts of calcein AM dye in MCF7-mCherry cells by flow cytometry presented as histogram plots.

**Immunogold electron microscopy**

We plated MCF7 monocultures or co-cultures of MCF7 and HS5 cells in 10 cm dishes at 12,000 cells per $cm^2$ with a 1:9 ratio of MCF7 to HS5 cells for co-cultures. After culturing cells for three days in LG/LF medium, we washed cells twice with PBS and then fixed cells with 2% paraformaldehyde and 0.2% glutaraldehyde (#15700 and #16200, respectively, Electron Microscopy Services, Hatfield, PA, USA) for one hour at 4°C. We then replaced the solution with ice-cold 4% paraformaldehyde with 0.1% Probumin® BSA (#820451, Millipore Sigma) in 200 mM HEPES and scraped the cells with a soft Teflon scraper (#07-200-366, Thermo Fisher). The frozen sample blocks were cryosectioned (60-70 nm thick section 10 μm apart) on EM grid. For immunolabelling, grids were washed with 2% gelatin (Cat# ES-006-B; Sigma-Aldrich) in phosphate buffer at 30-37 °C for 15-20 min followed by 10-20 mM (~0.15%) glycine for 5 min and 1% BSA/PBS for 2 min. Grids were incubated with 1:1 anti CX43 antibody (mouse mAb, epitope aa 247-382; BioLegend Cat#850001) and 1:10 anti GIV antibody (Cat#ABT80; Millipore) in 1% BSA/PBS for 1 hour at room temperature. For gold labelling the grids were incubated with Goat Anti-Mouse IgG tagged with 12 nm Colloidal Gold (Jackson Immuno research lab Cat#115-205-068) and Goat Anti-Rabbit IgG tagged with 6 nm Colloidal Gold (Jackson Immuno research lab Cat#111-195-144) diluted in 1:1 in 1% BSA for 1 h at room temperature followed by washing with PBS for 4 times. Then the grids were incubated with 1% Glutaraldehyde in PBS for 5 min. Then the grids were rinsed with distilled water 6 times. For contrast enhancement, the grids were incubated with 2% uranyl acetate at pH 7 for 5 min and in uranyl acetate (4%) (Cat# 21447-25; Polysciences) and methylcellulose (2%) (Cat# 9004-67-5; Sigma Aldrich) mixture for 5 min. Then the grids were dried in a wire loop at room temperature for 15 min. After that, the grids were mounted in Jeol 1400 plus Transmission Electron Microscope for visualization.

**Immunoblotting**

For immunoblotting in combination with GST pull-down assays (see below), we transfected Cos7 cells with Cx43-GFP (gift from David Spray (Addgene plasmid #69007; http://n2t.net/addgene:69007;RRID:Addgene_69007) (16)) using polyethylenimine (PEI) following the manufacturer's protocol as described in prior work (17, 18). Whole-cell lysates were prepared after washing cells with cold PBS before re-suspending and boiling them in sample buffer. For immunoblotting, protein samples were separated by SDS-PAGE and transferred to polyvinylidene fluoride membranes (Millipore Sigma). The membrane was stained with Ponceau S to visualize proteins Membranes were blocked with PBS supplemented with 5% nonfat milk before incubation with primary antibodies. Infrared imaging with dual-color detection and quantification was performed using a Li-Cor Odyssey imaging system. The anti-CX43 (BioLegend, CA, 247-382) antibody was used at 1:1000 v/v dilution. Figures were assembled for presentation using Adobe Photoshop and Illustrator. For co-culture experiments, we separated cancer cells from HS5 cells using EpCAM immunomagnetic beads as described above. We made whole-cell lysates using RIPA lysis buffer and quantified amounts of protein per sample by BCA assay (23235, Thermo Fisher Scientific). We used primary antibodies against GIV (H-6, Santa Cruz Biotechnology), CX43 (#3512, Cell Signaling Technology), or β-Actin (#4970, Cell Signaling Technology) and detected primary antibodies with appropriate anti-mouse or anti-rabbit secondary antibodies (#31430 and #31460, respectively, ThermoFisher) conjugated to horseradish peroxidase. We developed blots with ECL Plus Reagent (ThermoFisher) and quantified bands using ImageJ.

**GST pull-down assays**

Recombinant GST proteins were expressed in E. coli stain BL21 (DE3) and purified as previously described. Briefly, cultures were induced using 1mM IPTG overnight at 25°C. Cells were then pelleted and resuspended in either GST lysis buffer (25 mM Tris-HCl, pH 7.5, 20 mM NaCl, 1 mM EDTA, 20% (vol/vol) glycerol, 1% (vol/vol) Triton X-100, 2×protease inhibitor cocktail). Recombinant GST alone (control) or GST-tagged GIV proteins, GST-GIV-NT (1-166 aa) and GST-GIV-CT (1660-1870 aa) from *E. coli* were immobilized onto glutathione-Sepharose beads and incubated with binding buffer (50 mM Tris-HCl pH 7.4, 100 mM NaCl, 0.4% (v:v) Nonidet P-40, 10 mM $MgCl_2$, 5 mM EDTA, 2 mM DTT, 1X Complete protease inhibitor) overnight at 4°C temperature. For binding with cell lysates, cells were lysed in cell lysis buffer (20 mM HEPES pH 7.2, 5 mM Mg-acetate, 125 mM K-acetate, 0.4% Triton X-100, 1 mM DTT, 500 µM sodium orthovanadate, phosphatase inhibitor cocktail (Sigma-Aldrich, MO, USA) and protease inhibitor cocktail (Roche Life Science) using a 28G syringe, followed by centrifugation at 10,000Xg for 20 min. Cleared supernatant was then used in binding reaction with immobilized GST-proteins for 4 h at 4°C. After binding, bound complexes were washed four times with 1 ml phosphate wash buffer (4.3 mM $Na_2HPO_4$, 1.4 mM $KH_2PO_4$, pH 7.4, 137 mM NaCl, 2.7 mM KCl, 0.1% (v:v) Tween 20, 10 mM $MgCl_2$, 5 mM EDTA, 2 mM DTT, 0.5 mM sodium orthovanadate). Bound proteins were then eluted through boiling at 100°C in Laemmli sample buffer (BIORAD, CA, USA). The bound proteins were eluted at 37ºC for 10 min.

**Cell viability assays**

For viability assays, we seeded MCF7-CBG or MCF7-CBG-GIV cells at 10,000 cells per well in black wall 96 well plates (#165305, ThermoFisher Scientific). We washed cells once with PBS and then added various concentrations of tamoxifen, fulvestrant, or fulvestrant with 100 nM palbociclib (all from Cayman Chemical) in LG/LF medium with 4 wells per condition. Control wells received vehicle only. After three days in culture, we quantified relative numbers of viable cells by bioluminescence imaging using an IVIS Lumina system with Living Image 4.3.1 software (Perkin Elmer) as described previously (3). We plotted data as fraction change from vehicle only wells, which we defined as 1.

**Animal studies**

The University of Michigan Institutional Animal Care and Use Committee approved all experiments under protocol PRO00010534. We seeded MCF7-CBG or MCF7-CBG-GIV cells in T175 flasks. The following day, we changed medium to LG/LF medium without added estrogen and cultured cells overnight before harvesting cells with cell dissociation buffer (Gibco, ThermoFisher Scientific). We resuspended cells at 1 x $10^5$ cells per 100 µl sterile 0.9% NaCl for intracardiac injection. Prior to injection, we verified equal numbers of viable cells based on bioluminescence imaging of aliquots of each cell type. To generate systemic metastases, we injected 1 x $10^5$ MCF7-CBG or MCF7-CBG-GIV cells into the left ventricle of 7–10-week-old female NSG mice (strain NOD.Cg-*Prkdc*$^{scid}$ *Il2rg*$^{tm1Wjl}$/SzJ, #005557) (7-8 mice per group) (The Jackson Lab) as described previously by our lab (7). We performed bioluminescence imaging (IVIS Spectrum, Perkin Elmer) on mice at days indicated in the figure legend (7) We quantified total bioluminescence signal above background using Living Image software.

For orthotopic tumor xenografts of MDA-MB-231 breast cancer cells, we implanted 2.5 x $10^5$ parental or GIV KO MDA-MB-231 cells and 1 x $10^5$ human mammary fibroblasts into fourth mammary fat pads of 8-12-week-old female NSG mice (The Jackson Lab) as described previously (19).

**Statistics**

We performed all cell-based experiments at least three times and presented results as either a representative experiment or average of experiments ± SEM. For statistical analyses, we used python, R, GraphPad Prism software, or MATLAB (Mathworks). We assessed the statistical significance of Kaplan-Meier plots by log rank test. Kaplan-Meier analyses were performed using lifelines python package version 0.14.6. p-values for multiple comparisons were corrected using Tukey's method and repeated measure ANOVA was performed for time series and dose-dependent data.
